# Loss of *Mafb* and *Maf* distorts myeloid cell ratios and disrupts fetal mouse testis vascularization and organogenesis

**DOI:** 10.1101/2021.04.26.441488

**Authors:** Shu-Yun Li, Xiaowei Gu, Anna Heinrich, Emily G. Hurley, Blanche Capel, Tony DeFalco

**Author notes:** **Corresponding Author:** Tony DeFalco, **E-mail:**, **Address:** Division of Reproductive Sciences, Cincinnati Children’s Hospital Medical Center, 3333 Burnet Avenue, MLC 7045, Cincinnati, OH 45229 USA. These authors contributed equally to this work (Shu-Yun Li, Xiaowei Gu).

## Abstract

Testis differentiation is initiated when *Sry* in pre-Sertoli cells directs the gonad toward a male-specific fate. Sertoli cells are essential for testis development, but cell types within the interstitial compartment, such as immune and endothelial cells, are also critical for organ formation. Our previous work implicated macrophages in fetal testis morphogenesis, but little is known about genes underlying immune cell development during organogenesis. Here we examine the role of the immune-associated genes *Mafb* and *Maf* in mouse fetal gonad development, and we demonstrate that deletion of these genes leads to aberrant hematopoiesis manifested by supernumerary gonadal monocytes. *Mafb*;*Maf* double knockout embryos underwent initial gonadal sex determination normally, but exhibited testicular hypervascularization, testis cord formation defects, Leydig cell deficit, and a reduced number of germ cells. In general, *Mafb* and *Maf* alone were dispensable for gonad development; however, when both genes were deleted, we observed significant defects in testicular morphogenesis, indicating that *Mafb* and *Maf* work redundantly during testis differentiation. These results demonstrate previously unappreciated roles for *Mafb* and *Maf* in immune and vascular development and highlight the importance of interstitial cells in gonadal differentiation.

**Summary statement:** Deletion of *Mafb* and *Maf* genes leads to supernumerary monocytes in fetal mouse gonads, resulting in vascular, morphogenetic, and differentiation defects during testicular organogenesis.

## Introduction

Gonad morphogenesis is a highly orchestrated process involving germ cells, somatic supporting cells, interstitial/mesenchymal cells, immune cells, and vascular endothelial cells [1]. Cells in the fetal mouse testis undergo extensive cellular rearrangement between embryonic day (E) E11.5 and E12.5, which leads to the formation of testis cords, the basic structural unit of the testis. Testis cords are comprised of Sertoli and germ cells and give rise to seminiferous tubules in the adult organ [2]. Sertoli cells, the supporting cells of the testis that are the first sex-specific cell type specified in the XY gonad, express the sex-determining genes *Sry* and *Sox9* [3, 4]; in contrast, XX gonads are comprised of FOXL2+ granulosa cells, the female supporting counterparts of Sertoli cells. Sertoli cells are considered to be the critical cells that initiate testis cord morphogenesis and drive several aspects of testicular differentiation, including the specification of androgen-producing Leydig cells in the interstitial compartment [1]. However, other studies in the field suggest that endothelial and interstitial/mesenchymal cells play essential, active roles in testis morphogenesis, including driving testis cord formation and establishing a niche to maintain multipotent interstitial progenitor cells [5–10].

A growing body of evidence supports the idea that immune cells, such as macrophages, are critical players in organ formation and repair [11]. Other hematopoietic-derived cells, including myeloid cells like granulocytes and monocytes, also play a role in organ formation and homeostasis [12, 13], and infiltration by immune cells has been proposed to be a critical, fundamental part of the organogenesis program [14]. During development and growth, myeloid cells are implicated in the morphogenesis of multiple tissues, including bone, mammary gland ducts, heart, pancreatic islets, and retinal vasculature [15–19]. Within reproductive tissues, macrophages are critical for multiple aspects of ovulation, estrus cycle progression, and steroidogenesis in the adult ovary [20, 21], as well as Leydig cell development and spermatogonial stem/progenitor cell differentiation in the postnatal and adult testis, respectively [22, 23].

We have previously shown that depletion of macrophages, which represent the majority of immune cells in the early nascent gonad, disrupted fetal testicular vascularization and morphogenesis [8], demonstrating that immune cells are an integral part of the testicular organogenesis program. A recent single-cell study revealed there are multiple myeloid cell types in the mid-to-late-gestation fetal and perinatal testis, including monocytes and granulocytes [24]. However, the role of different immune cell types and how their numbers are regulated or balanced during organogenesis are still outstanding questions in the field. To carry out their diverse activities during organ development and function, macrophages and other myeloid cells have a diverse cellular repertoire, with potential to influence angiogenesis, tissue remodeling and clearing, and the production of various growth factors and cytokines [25, 26].

One group of proteins that may contribute to immune or macrophage function in the gonad is the family of large Maf bZIP transcription factors, which in mammals is comprised of MAFA, MAFB, and MAF (also called C-MAF). Mouse mutant models have demonstrated that *Mafb* and *Maf* play roles in different aspects of hematopoiesis, including macrophage differentiation within tissues and macrophage function during definitive erythropoiesis in the fetal liver [27–31]. In addition to regulating hematopoiesis, the large Maf factors influence cell fate decisions during the differentiation of multiple organs, including pancreas [32, 33], hindbrain [34, 35], eye [36], and kidney [29, 37]. The sole *Drosophila* ortholog of the mammalian large *Maf* genes, *traffic jam*, directs gonad development in flies via regulating cell adhesion molecules that mediate soma-germline interactions. Gametogenesis is stunted in *traffic jam* mutant testes and ovaries, leading to sterility in both sexes [38]. However, little is known about the functional role of large Maf factors in mammalian gonadal development.

In mice, MAFB and MAF, but not MAFA, are expressed in the fetal gonad [9, 39]. MAF is expressed in macrophages within the developing gonad-mesonephros complex [8], and MAFB has been well-characterized as a marker of monocyte-derived myeloid cells [29, 40]. In addition to immune cell expression, both MAFB and MAF are early markers of interstitial mesenchymal cells in both sexes [9], and in later stages of fetal development MAFB is expressed in both Leydig and Sertoli cells [39]. Knock-in mutant analyses and global conditional deletion of *Mafb* revealed that it was not required for fetal testicular differentiation or for maintenance of adult spermatogenesis [39]. However, multiple studies have suggested that *Mafb* and *Maf* have redundant or overlapping roles [27, 41], perhaps due to similar and conserved DNA binding domains [38, 42, 43]; additionally, *Mafb*;*Maf* double homozygous knockout embryos have earlier embryonic lethality than other combinations of large *Maf* genes [44], indicating that these genes have essential, redundant roles in embryogenesis.

Because of their previously reported roles in myeloid cell differentiation and in *Drosophila* gonad morphogenesis, here we have investigated the role of *Mafb* and *Maf* during mouse fetal gonad development. While *Mafb* or *Maf* alone were largely dispensable for fetal gonad differentiation, *Mafb*;*Maf* double-homozygous knockout embryos exhibited supernumerary monocytes, a newly-identified population of gonadal immune cells, that specifically localized near vasculature at the gonad-mesonephros border. Along with disrupted hematopoiesis in double-homozygous knockout embryos, we observed testicular hypervascularization, testis cord morphogenesis defects, and a reduction in germ cells in both sexes. Therefore, in conjunction with our previous findings, these results demonstrate that both reduced and increased numbers of immune cells disturb gonad differentiation. Additionally, double-homozygous knockout testes possessed a reduced number of Leydig cells. However, reduced Leydig cell differentiation in knockout testes was likely a secondary effect of hypervascularization driven by excess immune cells, as *Mafb*- and *Maf*-intact fetal testes in which vasculature was disrupted ex vivo also displayed reduced numbers of Leydig cells. In general, there appeared to be a stronger requirement for *Maf*, as compared to *Mafb*, in testicular differentiation; however, our data demonstrate that *Mafb* and *Maf* work redundantly to promote development of the testis by regulating differentiation of the interstitial compartment. This study provides evidence supporting the idea that immune cell activity and number are delicately balanced during organogenesis to regulate vascular and tissue remodeling processes that are integral to testis morphogenesis.

## Materials and methods

### Mice

All mice used in this study were housed in the Cincinnati Children’s Hospital Medical Center’s or Duke University Medical Center’s animal care facility, in compliance with institutional and National Institutes of Health guidelines. Ethical approval was obtained for all animal experiments by the Institutional Animal Care and Use Committee (IACUC) of Cincinnati Children’s Hospital Medical Center or Duke University Medical Center. Mice were housed in a 12-hour light/dark cycle and had access to autoclaved rodent Lab Diet 5010 (Purina, St. Louis, MO, USA) and ultraviolet light-sterilized RO/DI constant circulation water ad libitum.

CD-1 (Charles River stock #022) and C57BL/6J (Jackson Laboratory stock #000664) mice were used for wild-type expression studies. *Mafb^GFP^* (*Mafb*^tm1Jeng^), a null *Mafb* knock-in allele in which the eGFP coding sequence replaced the *Mafb* coding sequence [29], was a gift from S. Takahashi (University of Tsukuba, Japan) via L. Goodrich (Harvard University). *Maf^-/-^* knockout mice (*Maf*^tm1Glm^) [45] were a gift from I.C. Ho (Harvard University/Brigham and Women’s Hospital). *Mafb* and *Maf* knockout mice were maintained and bred as double-heterozygous animals, which were on a mixed background of BALB/c, C57Bl/6J, and CD-1. *Vav1*-Cre mice (Tg(Vav1-cre)1Graf), which target all cells of the hematopoietic lineage [46], were a gift from H.L. Grimes. *Lyz2*-Cre mice (*Lyz2^tm1(cre)Ifo^*; also called *LysM*-Cre), which target myeloid cells [47], were purchased from Jackson Laboratories (stock #004781). *Csf1r*-Cre mice (Tg(Csf1r-icre)1Jwp) (Jackson Laboratory stock #021024), which target monocytes and macrophages [48], were a gift from R. Lang (Cincinnati Children’s Hospital). *Rosa*-Tomato mice (*Gt(ROSA)26Sor^tm14(CAG-tdTomato)Hze^*), used as a Cre-responsive fluorescent reporter strain [49], were purchased from Jackson Laboratory (stock #007914). *Cx3cr1*-GFP mice *(Cx3cr1^tm1Litt^*), a knock-in *Cx3Cr1* reporter allele which is expressed in monocytes and differentiated tissue-resident macrophages, subsets of NK and dendritic cells, and brain microglia [50], were purchased from Jackson Laboratory (stock #005582). *Ccr2*-GFP mice (*Ccr2^tm1.1Cln^*) [51], in which infiltrating monocytes are labeled with GFP, were purchased from Jackson Laboratory (stock #027619).

Presence of the *Mafb^GFP^* knock-in allele was assessed via a fluorescent dissecting microscope to visualize GFP fluorescence in embryos, particularly in the central nervous system, snout, eye lens, and interdigital webbing, and was confirmed by PCR genotyping for GFP with primers 5’ GAC GTA AAC GGC CAC AAG TT 3’ and 5’ AAG TCG GTG CTG CTT CAT GTG 3’. Presence of the *Mafb* wild-type allele was determined via PCR genotyping with primers 5’ GGT TCA TCT GCT GGT AGT TGC 3’ and 5’ GAC CTT CTC AAG TTC GAC GTG 3’. The *Maf* knockout allele was identified by PCR genotyping with primers 5’ TGC TCC TGC CGA GAA AGT ATC CAT CAT GGC 3’ and 5’ CGC CAA GCT CTT CAG CAA TAT CAC GGG TAG 3’ specific to the neomycin cassette in the knockout allele. Presence of the *Maf* wild-type allele was determined by presence of immunofluorescent MAF staining in the eye lens and in eye-associated macrophages, using a whole-mount immunofluorescence protocol (described below) on dissected fetal eyes. All PCR genotyping was performed on tail DNA isolated via an alkaline lysis protocol.

To obtain fetal samples at specific stages, timed matings were arranged in which a single adult male was paired with one or two adult females. Noon on the day a vaginal plug was observed was designated as E0.5.

Throughout the manuscript, we define the “control” genotype as: *Mafb*^+/+^ or *Mafb^GFP/+^* for *Mafb* experiments; *Maf*^+/+^ or *Maf^+/-^* for *Maf* experiments; and *Maf^+/-^*; *Mafb^GFP/+^* for compound heterozygous+knockout (compound heterozygous+KO, defined as 3 of 4 copies of *Mafb* and *Maf* are KO alleles) experiments and double knockout (double KO, defined as all 4 copies of *Mafb* and *Maf* are KO alleles) experiments. We define the “*Mafb* single KO” genotype as *Mafb^GFP/GFP^*; the “*Maf* single KO” genotype as *Maf^-/-^*; and the “double KO” genotype as *Mafb^GFP/GFP^*;*Maf^-/-^*. For compound heterozygous+KO analyses, we define the “*Mafb* KO;*Maf-*heterozygous” genotype as *Mafb^GFP/GFP^*;*Maf^+/-^* and the “*Mafb*-heterozygous;*Maf* KO” genotype as *Mafb^GFP/+^*;*Maf^-/-^*.

### Immunofluorescence

Whole-mount immunofluorescence was performed as previously described [10]. Gonads were dissected in phosphate-buffered saline (PBS) and fixed overnight at 4°C in 4% paraformaldehyde (PFA) with 0.1% Triton X-100. After several washes in PBTx (PBS + 0.1% Triton X-100), samples were incubated in a blocking solution (PBTx + 10% fetal bovine serum [FBS] + 3% bovine serum albumin [BSA]) for 1–2 hours at room temperature. Primary antibodies were diluted in blocking solution and samples were rocked in primary antibodies overnight at 4°C. After several washes in PBTx, fluorescent secondary antibodies were applied for 3–5 hours rocking at room temperature or overnight at 4°C. After several washes in PBTx, samples were mounted on slides in Fluoromount-G (SouthernBiotech) or 2.5% DABCO (Sigma-Aldrich) in 90% glycerol. For co-labeling of anti-CD11b or anti-F4/80 antibody with anti-CD45 antibody (which are all raised in rat), a sequential stain was done, in which F4/80 or CD11b was first incubated and labeled with fluorescently-conjugated Cy3 anti-rat secondary followed by extensive washes and subsequent incubation with anti-CD45 antibody directly conjugated with Alexa Fluor 488. Primary antibodies used are listed in Supplemental Table S2. Alexa-488, -555, -568, and -647-conjugated secondary antibodies (Molecular Probes) were all used at 1:500. Dy-Lite 488 donkey anti-chicken and Cy3 donkey anti-rat secondary antibodies (Jackson Immunoresearch) were used at 1:500. Nuclei were stained with 2 μg/ml Hoechst 33342 (Molecular Probes/Life Technologies/Thermo Fisher) or DAPI (Sigma-Aldrich). Samples were imaged either on a Nikon Eclipse TE2000 microscope (Nikon Instruments) with an Opti-Grid structured illumination imaging system using Volocity software (PerkinElmer, Waltham, MA, USA), a Nikon A1 inverted confocal microscope (Nikon Instruments), or a Leica SP2 confocal microscope (Leica Microsystems).

### Germ cell depletion

Pregnant CD-1 females were injected intraperitoneally at E10.5 with 100 μl of 16 mg/ml busulfan (Sigma-Aldrich) dissolved in 50% DMSO or an equivalent volume of 50% DMSO, as previously described [52]. Embryos were harvested at E13.5 and processed for immunofluorescence.

### RNA extraction, cDNA synthesis, and quantitative real-time PCR (qRT-PCR)

Total RNA was extracted and processed for quantitative real-time PCR (qRT-PCR). Tissue was homogenized in 200µl TRIzol reagent (Invitrogen/Thermo Fisher). RNA extraction was then performed using a TRIzol/isopropanol precipitation method. Briefly, 40 µL of chloroform was added to the TRIzol/tissue mixture, shaken by hand, incubated at room temperature for 3 minutes, and centrifuged at 12,000 × g for 10 minutes at 4°C. The upper aqueous layer was carefully recovered and added to 80 µL isopropanol and 0.4 µL GlycoBlue coprecipitant (Thermo Fisher Scientific), which was rocked at room temperature for 10 minutes. After centrifugation at 12,000 × g for 10 minutes at 4°C, supernatant was removed, and the pellet was washed with 500 µL of ethanol. After another centrifugation (with same parameters), the RNA pellet was briefly air-dried and diluted in nuclease-free water. RNA quality was assessed by spectrophotometric analysis via absorbance at 260 and 280 nm, in which only RNA samples with a 260/280 ratio greater than or equal to 1.6 was used for qRT-PCR analysis (although sample ratios were usually between 1.7-2.0). An iScript cDNA synthesis kit (BioRad) was used on 500ng of RNA for cDNA synthesis. qRT-PCR was performed using the Fast SYBR Green Master Mix (Applied Biosystems/Thermo Fisher) on the StepOnePlus Real-Time PCR system (Applied Biosystems/Thermo Fisher). The following parameters were used: 95°C for 20s, followed by 40 cycles of 95°C for 3s and 60°C for 30s, followed by a melt curve run. Primer specificity for a single amplicon was verified by melt curve analysis. *Gapdh* was used as an internal normalization control.

### Ex vivo whole gonad droplet culture

Whole gonad-mesonephros complexes from E12.5 male CD-1 embryos were dissected in PBS and cultured for 48 hours at 37 °C and 5% CO_2_ in 30 µl droplets containing DMEM medium with 5% (or 10% in PDGF-BB + VEGFR-TKI II experiments) fetal bovine serum (FBS) (for VEGFA_165_ and PDGF-BB alone experiments) and 1% penicillin–streptomycin, as described previously [10, 53]. For PDGF-BB experiments, recombinant rat PDGF-BB (R&D Biosystems #520-BB, 50 ng/mL) or equivalent amount of 0.1% BSA vehicle was added to media. For VEGFA experiments, recombinant murine VEGFA_165_ (PeproTech #450-32, 50 ng/mL) or equivalent amount of 0.1% BSA vehicle was added to media. For VEGFR-TKI II experiments, VEGF Receptor Tyrosine Kinase Inhibitor II (VEGFR-TKI II; Calbiochem/EMD Millipore #676481-5MG, 1.8 μg/mL) or equivalent amount of DMSO vehicle was added to media.

For PDGF-BB experiments in Figure 8, 5% FBS media was used since the baseline amount of vasculature is lower and hypervascularization can be more easily induced upon PDGF-BB treatment. Thus, upon this increase in vasculature, there is a visible reduction of Leydig cell number relative to controls in these conditions. To address whether the reduction of Leydig cells in the above experiment was caused by hypervascularization or is a direct negative effect of PDGF-BB treatment on Leydig cell differentiation, in Supplemental Figure 8 we used 10% FBS media, which has a higher baseline amount of vasculature relative to 5% FBS (as seen in Figure 8), so we can block the hypervascularization caused by 10% FBS (via additional simultaneous treatment with VEGFR-TKI II) to determine more definitively if PDGF-BB has any direct negative effect on Leydig cell number in the absence of hypervascularization.

After culture, gonads were fixed in 4% PFA for immunofluorescence and processed for whole-mount immunofluorescence as described above; alternatively, gonads were separated from the mesonephros for RNA extraction and qRT-PCR analysis as described above.

### Microarray Analysis

Purified populations of E12.5 XY *Mafb*-GFP-positive cells were obtained from three independent pairs of *Mafb^GFP/+^*;*Maf^+/-^* testes (control) and three independent pairs of *Mafb^GFP/+^*;*Maf^-/-^* (mutant) testes via FACS as previously described [54]. The testis was left attached to the gonad/mesonephros border region and its associated macrophage and interstitial cell populations [9]. Total RNA was extracted from approximately 100,000 GFP-positive cells per biological replicate using the an RNeasy Micro Kit (Qiagen), with modifications as previously described [54], and submitted to the Duke University Microarray Facility for labeling and hybridization to Affymetrix GeneChip Mouse Genome 430A 2.0 microarrays. Data analysis was performed with Affymetrix Expression Console Software using an RMA (Robust Multi-Array Average) algorithm and transformed into log base 2. Genes that had 1.5-fold-or-higher fold change with a *P-*value of less than 0.05 were considered significantly upregulated or downregulated. The raw data is available at the Gene Expression Omnibus (GEO) under accession number GSE41715.

### Germ cell quantification and testis cord morphometric analyses

Germ cells of E11.5 XY gonads were labeled by anti-SOX2 antibody and testis cords of E13.5 XY gonads were visualized by anti-AMH antibody. For meiotic germ cell counts, the number of SYCP3+ germ cells were counted per total germ cells, as marked by PECAM1 or CDH1. For all quantifications, a sample size of *n*=3-10 gonads for each genotype were analyzed using ImageJ software (NIH). For E11.5 XY gonads, the number of SOX2+ germ cells per optical section (within a field of view 375 μm wide) were counted manually; three separate optical sections of each gonad were counted and averaged. For E13.5 XY gonads, five testis cords of each gonad in each image (within a field of view 750 μm wide) were measured and averaged. Surface-biased longitudinal optical sections that showed the full height of the cords were used for height measurements, while both longitudinal and transverse sections of cords were used for width measurements. For E13.5 XX gonads, three or four separate optical sections per gonad were analyzed and averaged for both total germ number and SYCP3+ cell number.

### Sample sizes and statistical analyses

For qRT-PCR, fold change in mRNA levels was calculated relative to controls using a ΔΔC_t_ method. Results were shown as mean ± SD. An unpaired, two-tailed Student t-test was performed to calculate *P* values based on ΔC_t_ values, in which *P*<0.05 was considered statistically significant. Statistical analyses were performed using Prism version 5.0 (GraphPad). At least three gonads from independent embryos (*n*≥3) were used for qRT-PCR analyses. For ex vivo gonad culture, at least three independent experiments were performed and within each experiment at least 3 gonads from independent embryos (*n*≥3) were pooled for each biological replicate. For immunofluorescence assays, at least three independent experiments were performed and within each experiment multiple gonads from independent samples (*n*≥2) were used. For germ cell quantifications and morphometric analyses, sample sizes are listed above for each group. Data is represented as mean ± SD, and statistical significance was determined by an unpaired, two-tailed Student t-test in which *P*<0.05 was considered statistically significant.

## Results

### Initial gonadal sex differentiation occurs normally in the absence of Mafb or Maf

As a first step to investigate the role of the large Maf factors, we examined basic aspects of male and female gonadal sex differentiation in *Mafb* single KO or *Maf* single KO gonads. Relative to control XY littermates, we observed comparable numbers of Sertoli cells in XY *Mafb* single KO and *Maf* single KO fetal gonads (Supplemental Figure S1A-C). While we did note some testis cord formation defects and reduced germ cell numbers in E12.5 *Mafb* single KO and *Maf* single KO testes (Supplemental Figure S1D-F), testis cord structure and germ cell numbers generally recovered by E13.5 (Supplemental Figure S1G-I). We also observed that there were comparable numbers of Leydig cells in E13.5 control versus *Mafb* single KO and *Maf* single KO testes (Supplemental Figure S1J-L). However, in E13.5 *Maf* single KO testes, we noticed subtle defects in cord architecture and some disruptions of the surface coelomic artery [55], in which the vessel was disorganized and multi-layered (Supplemental Figure S1M-O). These data indicate that male gonadal sex differentiation generally occurred normally in *Mafb* single KO or *Maf* single KO testes, but we did note that *Maf* single KO mutant testes were smaller than controls.

We observed FOXL2+ cells throughout E13.5 XX *Mafb* single KO and *Maf* single KO fetal gonads comparably to control XX gonads (Supplemental Figure S2A-C), indicative of ovarian differentiation. Another aspect of fetal ovarian differentiation is the entry of germ cells into meiosis, marked by SYCP3 (synaptonemal complex protein 3) expression, starting at E13.5, which does not occur in the fetal testis [56]. As in E14.5 control XX gonads (Supplemental Figure S2D), germ cells throughout E14.5 XX *Mafb* single KO and *Maf* single KO gonads expressed SYCP3 (Supplemental Figure S2E and F). These findings demonstrate that female gonadal sex differentiation and germ cell differentiation occurred normally in the absence of *Mafb* or *Maf.* However, as with fetal testes, we observed that *Maf* single KO mutant ovaries were smaller than their control counterparts. Overall, our data indicate that initial gonadal sexual differentiation in either sex does not require *Mafb* or *Maf*.

### Double KO gonads undergo initial gonadal sex differentiation normally

To address the possibility that *Mafb* and *Maf* act redundantly during gonad development, we examined compound heterozygous+KO and double KO embryos. Adult double-heterozygous control males and females were fertile and able to generate double KO embryos. However, since double KO embryos only survive until E13.5 [44], we focused on early aspects of gonad differentiation.

First, to assess if initial gonadal sex differentiation and supporting cell specification occurred normally in double KO gonads, we examined the expression of SOX9 and FOXL2, markers for Sertoli and pre-granulosa cells, respectively. We found that E13.5 XY double KO gonads expressed SOX9 within Sertoli cells, similar to controls (Figure 1A and B). However, mutant testes were smaller than controls. E13.5 XX double KO gonads expressed FOXL2 within pre-granulosa cells, similar to controls (Figure 1C and D), but, as with testes, double KO ovaries were smaller than their control counterparts. Overall, these data indicate that initial gonadal sex differentiation occurred normally in the absence of both *Mafb* and *Maf*.

**Figure 1.**
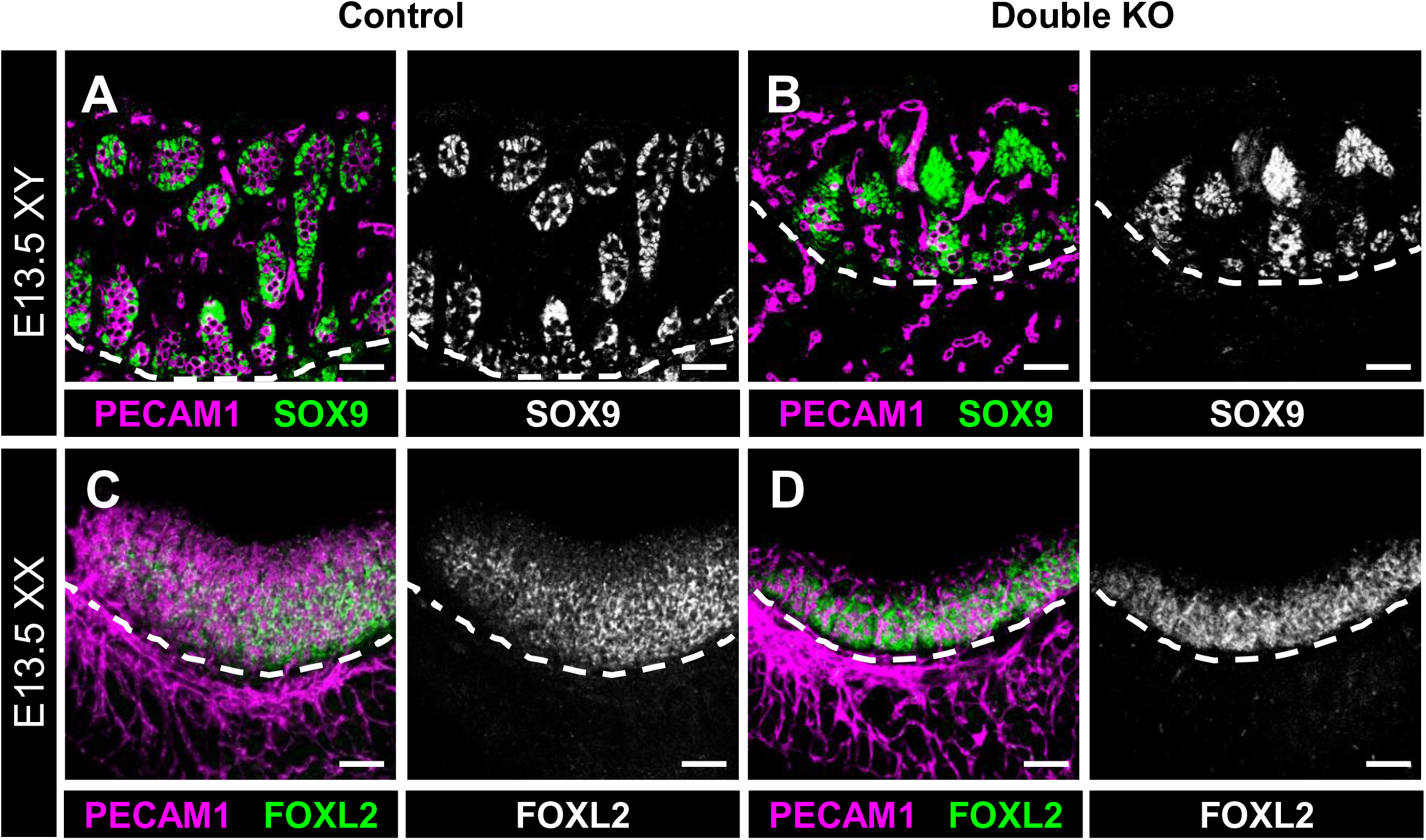
Initial gonadal sex differentiation occurs normally in double KO gonads. Immunofluorescent images of E13.5 XY (A,B) and XX (C,D) control (A,C) and double KO (B,D) fetal gonads. White dashed lines indicate gonad-mesonephros border. Gonadal sex differentiation, as assessed by presence of SOX9+ Sertoli cells in XY gonads (A and B) and FOXL2+ pre-granulosa cells in XX gonads (C and D), appears to develop normally in both control (A and C) and double KO (B and D) gonads. Scale bars, 100 μm.

### Cord morphogenesis is disrupted and germ cells are reduced in double KO testes

We next examined testis cord formation in E13.5 XY compound heterozygous+KO and double KO gonads relative to control littermate gonads. In general, testis cords were smaller in *Mafb-*heterozygous;*Maf* KO and double KO testes (Figure 2A-D), likely due to decreased numbers of germ cells. *Mafb* KO;*Maf*-heterozygous testes appeared grossly normal in their cord architecture (Figure 2B). However, in *Mafb-*heterozygous;*Maf* KO samples, cord abnormalities such as fused or branched cords were more common than in controls (Figure 2C), and we noted a more severe germ cell deficit as compared to *Mafb* KO;*Maf*-heterozygous mutant testes. In double KO testes, there were more dramatic perturbations in testis cord structure as compared to other genotypes (Figure 2D). While Sertoli cells were aggregated and usually sorted out from interstitial cells, virtually all cords were fused or branched, resulting in dramatic disorganization of cords in double KO testes. Morphometric analyses confirmed a reduction in testis cord height and width in E13.5 XY compound heterozygous+KO and double KO gonads (Figure 2M and N). Overall, our analyses showed that *Maf* plays a more critical role than *Mafb* in testis cord formation, but double KO gonads generally had the most severe phenotype.

**Figure 2.**
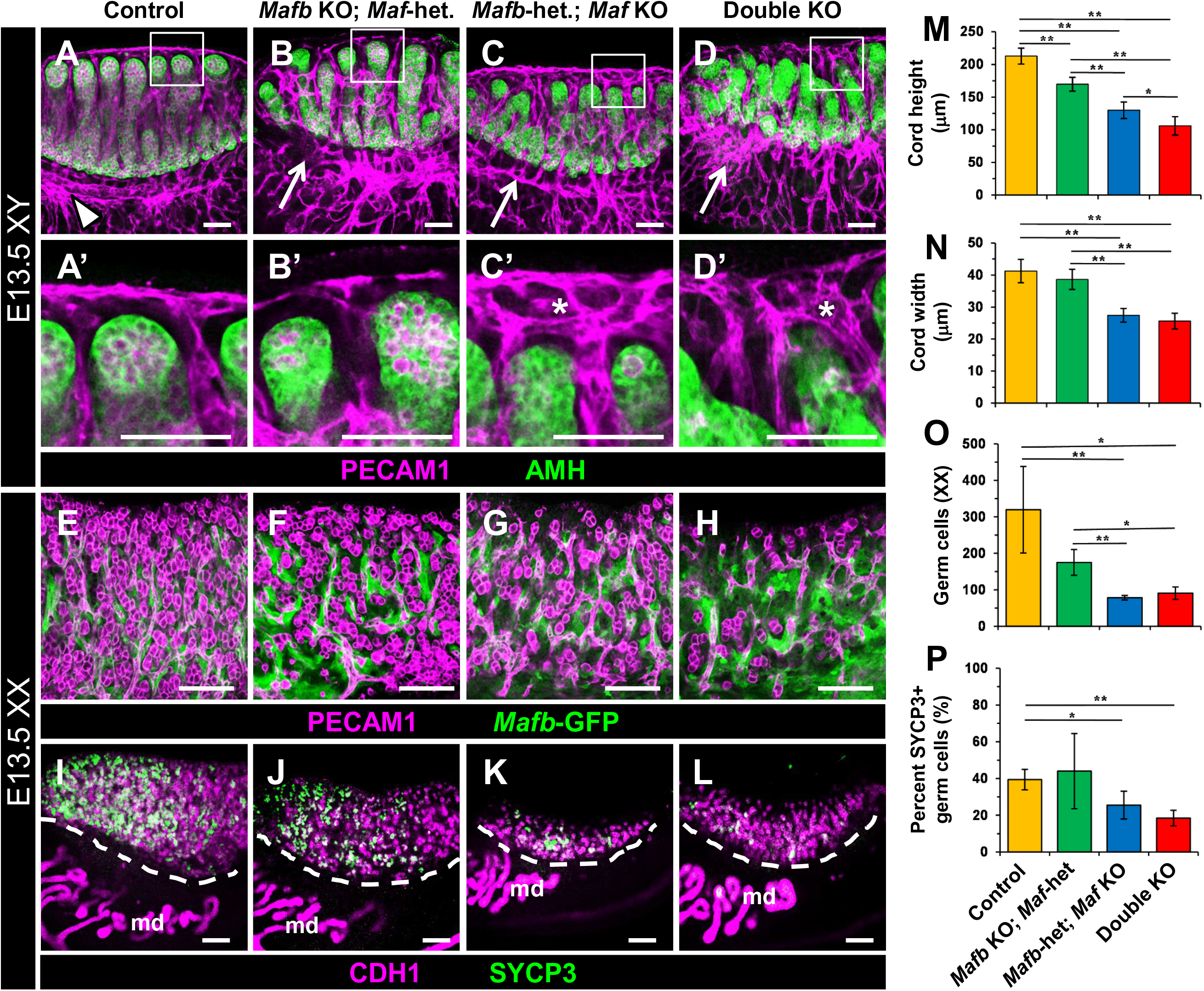
*Mafb* and *Maf* act redundantly to regulate gonad differentiation. Immunofluorescent images of E13.5 XY (A-D) and E13.5 XX (E-L) control (A, E, I), *Mafb* KO;*Maf*-heterozygous (B, F, J), *Mafb-*heterozygous;*Maf* KO (C, G, K), and double KO (D, H, L) gonads. A’-D’ are higher-magnification images of the boxed regions of coelomic vessels in A-D. Control (A) testes contained a single-vessel coelomic artery and robust, well-defined vascular plexus at the gonad-mesonephros border (A, arrowhead). In *Mafb* KO;*Maf*-heterozygous (B), *Mafb-*heterozygous;*Maf* KO (C) and double KO (D) gonads, the vascular plexus was disorganized (B-D, arrows). *Mafb-*heterozygous;*Maf* KO (C, C’) and double KO (D, D’) gonads also had extensive hypervascularization in the region of the coelomic vessel (asterisks in C’ and D’). (E-I) Compared to E13.5 control (E and I) ovaries, *Mafb-*heterozygous;*Maf* KO (G and K) and double KO (H and L) contained fewer SYCP3+ meiotic germ cells. *Mafb-*heterozygous;*Maf* KO (G and K) and double KO (H and L) ovaries also had an overall reduced number of PECAM1+/CDH1+ germ cells. md, mesonephric ducts. Scale bars, 100 μm. (M-P) Graphs showing quantification of testis cord height in E13.5 XY gonads (M), testis cord width in E13.5 XY gonads (N), number of total germ cells per optical section in E13.5 XX gonads (O), and percentage of germ cells expressing SYCP3 in E13.5 XX gonads (P). All graph data are represented as mean +/- SD. *, *P*<0.05; **, *P*<0.01 (Student t-test).

Time course analysis of compound heterozygous+KO gonads revealed that germ cells were present in numbers comparable to controls at E9.5, but germ cells in KO embryos were aberrantly localized outside the hindgut while migrating and were consequently dramatically reduced in number by E10.5 and E11.5, particularly in *Mafb*-heterozygous;*Maf* KO gonads (Figure 3A-I). These data suggest that defective germ cell migration is an underlying cause of germ cell reduction in KO gonads. We found that germ cells themselves do not express MAFB or MAF (Figure 3J-L), indicating that germ cell reductions were likely not due to germ-cell-intrinsic defects. We did not observe any defects in cell cycle status or proliferation, nor an increase in apoptosis, in germ cells in double KO gonads (Supplemental Figure S3A-H). We also investigated expression levels of *Cxcl12* (also called *sdf-1*) and *Kitl* (*kit ligand*; also called *stem cell factor*), which encode critical ligands secreted by the soma to recruit germ cells to the gonad, but did not find any significant changes in *Cxcl12* or *Kitl* mRNA expression within E11.5 XY *Mafb*-heterozygous;*Maf* KO gonad/mesonephros complexes relative to controls (Supplemental Figure S3I). Finally, to address if any defects in gonad size and germ cell colonization were due to disruptions in initial sex determination, we examined the expression of *Sox9*, a testis-specific gene, and *Foxl2* and *Wnt4*, ovary-specific genes, in E13.5 XX and XY *Mafb*-heterozygous;*Maf* KO gonads via qRT-PCR, and we found no significant changes in expression (Supplemental Figure S3J).

**Figure 3.**
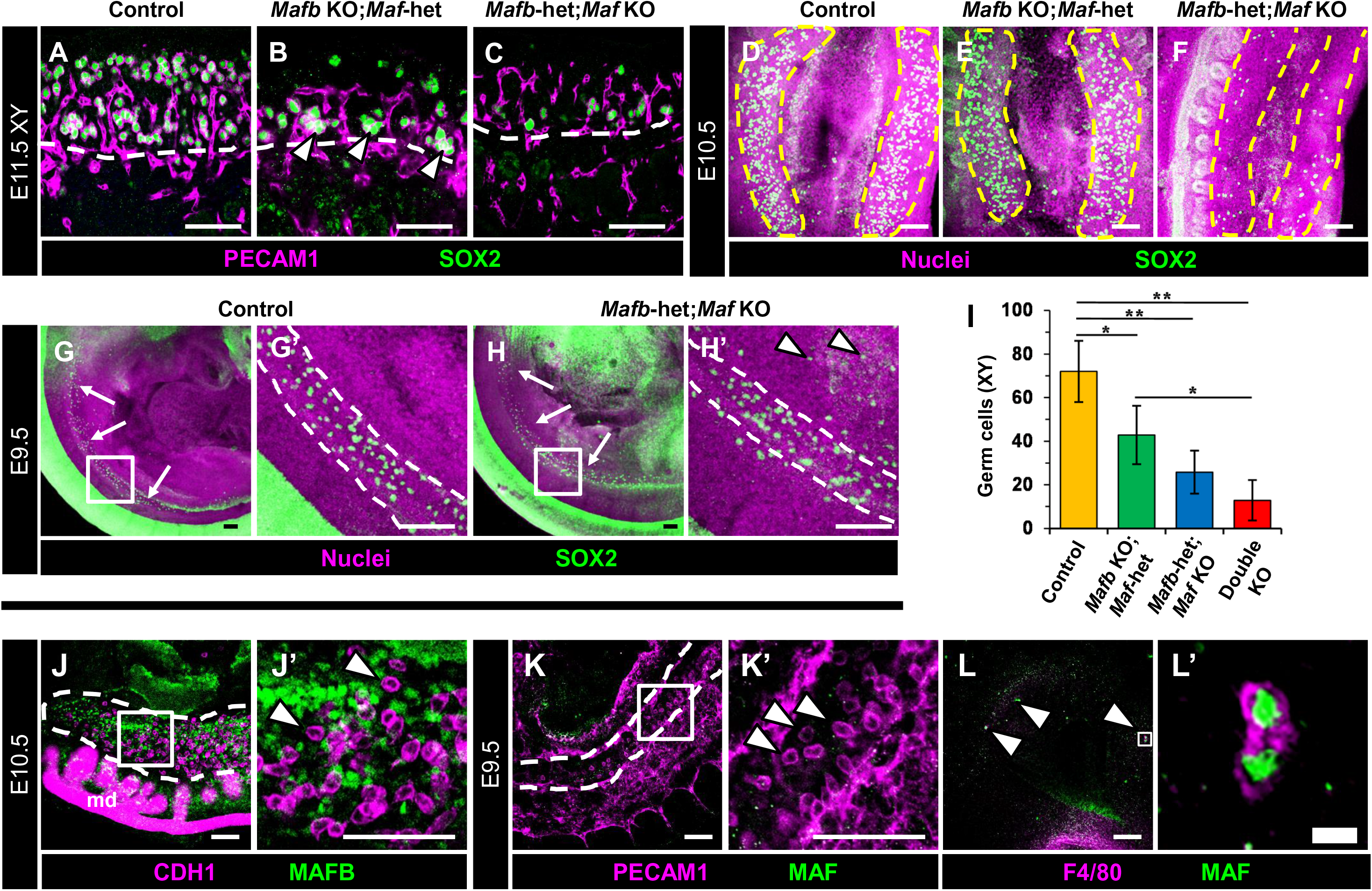
Germ cell colonization is disrupted in *Mafb*-heterozygous;*Maf* KO gonads. (A-H) Immunofluorescent images of E11.5 XY gonads (A-C), E10.5 urogenital ridges (D-F), and E9.5 embryonic trunks (G and H) from control (A,D,G), *Mafb* KO;*Maf*-heterozygous (B,E), and *Mafb*-heterozygous;*Maf* KO (C,F,H) embryos. White dashed lines indicate gonad-mesonephros border in A-C and hindgut boundaries in G and H; yellow lines mark gonad boundaries in D-F. G’ and H’ are higher-magnification images of the boxed regions in G and H. E11.5 XY *Mafb* KO;*Maf*-heterozygous gonads show enrichment of germ cells at the gonad-mesonephric border (arrowheads in B), while *Mafb*-heterozygous;*Maf* KO gonads (C) exhibit a significant overall reduction in germ cells at E10.5 and E11.5 (C and F). At E9.5, control (G) and *Mafb*-heterozygous;*Maf* KO embryos (H) show similar numbers of germ cells along the embryo axis (arrows in G and H), but germ cells are aberrantly localized outside the hindgut in *Mafb*-heterozygous;*Maf* KO embryos (arrowheads in H’). (I) Graph showing quantification of germ cells per optical section in E11.5 XY gonads of different genotypes. Data are represented as mean +/- SD. *, *P*<0.05; **, *P*<0.01 (Student t-test). (J-L) Immunofluorescence images of E10.5 urogenital ridge (J), E9.5 embryonic trunk (K), and E9.5 head (L) from wild-type CD-1 embryos. Dashed outline marks gonad boundary in J and hindgut boundaries in K. J’-L’ are higher-magnification images of the boxed regions in J-L. MAFB is expressed in gonadal and mesonephric mesenchyme (arrowheads in J’) but not in CDH1+ germ cells. MAF is not expressed in E9.5 PECAM1+ germ cells (arrowheads in K’) migrating in the hindgut, but it is expressed in F4/80+ macrophages in the E9.5 head (arrowheads in L). md, mesonephric ducts. Thin scale bars, 100 μm, thick scale bar, 10 μm.

One potential concern was whether the lack of germ cells in XY KO gonads affected their ability to form testis cords efficiently. To investigate this possibility, we examined the development of fetal testes in which germ cells were ablated. Using busulfan administration in utero to efficiently deplete germ cells from the fetal testis (Supplemental Figure S4A and B), we assessed effects on testis morphogenesis. Germ cell ablation revealed no gross differences in Sertoli cell specification, testis cord structure, or vascular development between germ-cell-depleted testes and vehicle control testes at E13.5 (Supplemental Figure S4A-D). These data suggest that *Mafb* and *Maf* affect testis cord structure independently of their effects on germ cell numbers, and that the two genes act redundantly to promote proper testis cord formation.

### Double KO ovaries exhibit germ cell reduction with a potential delay in meiosis onset

In E13.5 XX KO gonads, similarly to E13.5 XY KO gonads, we found progressively more severe phenotypes as we knocked out more copies of *Mafb* and *Maf*, whereby *Maf* mutation had more deleterious effects than *Mafb* mutation. We found that XX double KO gonads were smaller, with a reduction in germ cells (Figure 2E-H and O), although *Mafb-*heterozygous;*Maf* KO mutant gonads were occasionally similar or more severe in germ cell loss as compared to *Mafb* KO;*Maf*-heterozygous gonads (Figure 2E-L and O). To investigate the onset of meiotic entry as a readout of ovarian differentiation, we examined the expression of SYCP3. While there were widespread SYCP3+ cells in the anterior of E13.5 control ovaries, there were many fewer SYCP3+ cells in compound heterozygous+KO and double KO ovaries (Figure 2I-L). Quantification of total germ cells and SYCP3+ cells revealed that E13.5 *Mafb*-heterozygous;*Maf* KO and double KO ovaries showed not only a reduction in total germ cell number, but also a reduced percentage of germ cells expressing SYCP3 (Figure 2O and P). These data suggest that there was a delay in meiotic entry in *Mafb*-heterozygous;*Maf* KO and double KO gonads. It is possible, and perhaps likely, that they could recover later in development, as we observed in single KO gonads, but we could not address this possibility due to early embryonic lethality of double KO embryos after E13.5.

### Vascular remodeling is severely disrupted in double KO testes

Vascularization is a hallmark of testicular differentiation that is critical for testis cord morphogenesis and maintenance of Leydig progenitor cells [5–7, 10], and is characterized by formation of a main coelomic artery and a vascular plexus that runs along the gonad-mesonephros border [55, 57]. In E13.5 control and *Mafb* KO;*Maf*-heterozygous testes, a coelomic artery was visible (Figure 2A and B), although there were some disruptions in the gonad-mesonephric vascular plexus in *Mafb* KO;*Maf*-heterozygous testes (Figure 2B). In *Mafb-*heterozygous;*Maf* KO testes, the characteristic coelomic artery was still present, but was disorganized and multi-layered rather than structured as a single vessel; additionally, the vascular plexus at the mesonephric border was disorganized and thinner relative to control samples (Figure 2C). This phenotype was similar to, but more severe than, *Maf* single KO testes (Supplemental Figure S1O).

In double KO testes, the vascular phenotype was even more dramatic, and was characterized by extensive hypervascularization throughout the gonad. Instead of a well-defined vascular plexus and an avascular mesonephric region adjacent to the gonad as in controls (Figure 4A), in double KO testes endothelial cells were highly disorganized, resulting in a severe disruption of the mesonephric vascular plexus (Figure 4B). Additionally, the coelomic artery was multi-layered and severely disorganized as compared to controls. Therefore, for vascularization, *Maf* appeared to be more critical than *Mafb*, but, ultimately, mutating all copies of *Mafb* and *Maf* resulted in the most severe defects.

**Figure 4.**
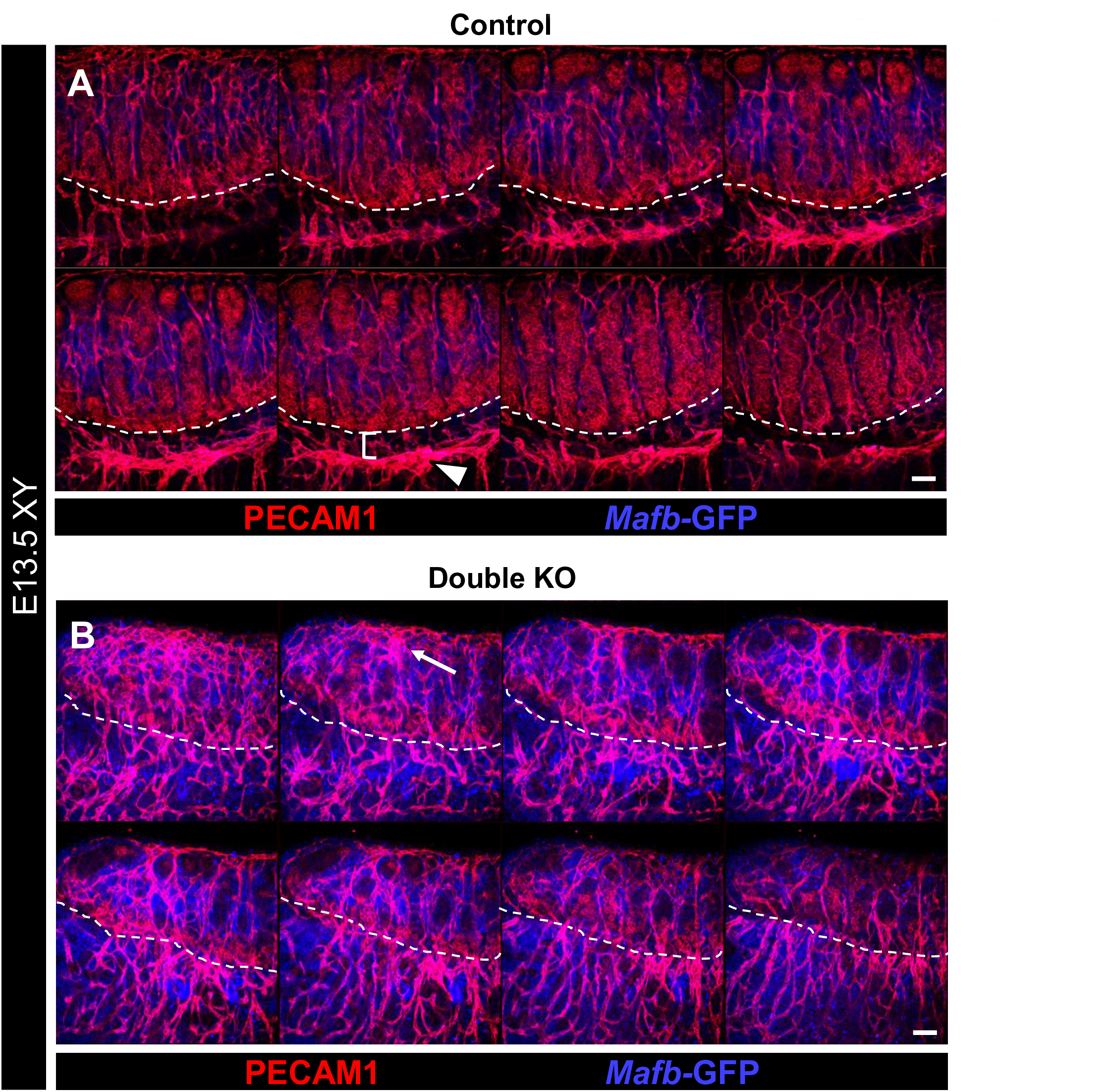
Double KO testes exhibit disruptions in vascular patterning. Immunofluorescent images of E13.5 control (A) and double KO (B) fetal testes. Each panel shows 8 consecutive confocal optical sections equally spaced through the entire gonad. Dashed lines indicate gonad-mesonephros boundary. Control testes (A) possess a fully developed gonad-mesonephric vascular plexus (arrowhead in A) with a well-defined avascular region between the vascular plexus and gonad (bracket in A). In contrast, double KO gonads (B) exhibit severely disrupted vascular development, with a hypervascularized coelomic surface (arrow) and aberrantly vascularized gonad-mesonephric border region. Scale bars, 100 μm.

### Maf KO gonad/mesonephros complexes exhibit ectopic CD11b-bright immune cells

In our analyses of KO gonads, we observed changes in the GFP expression pattern (from the *Mafb^GFP^* allele) in *Maf* KO gonads. In addition to the interstitial mesenchyme GFP expression we previously reported [9], in *Maf* KO gonads there were numerous ectopic GFP-bright round cells scattered throughout the gonad and mesonephros, although mostly concentrated in the mesonephros near the highly vascularized gonadal border, in both fetal testes and ovaries (Supplemental Figure S5A-F). The shape and localization of the ectopic GFP-positive cells within KO gonads, in addition to previous reports of *Mafb^GFP^* expression in macrophages [29], suggested that these cells were immune cells. Therefore, we investigated whether loss of *Mafb* or *Maf* function affected hematopoietic cells in the gonad/mesonephros complex. We first examined F4/80-positive macrophages, a prevalent immune cell in the fetal gonad, but detected no differences in F4/80 expression in *Mafb* single KO or *Maf* single KO gonads relative to controls (Figure 5A-D). In contrast, there was a dramatic change in the pattern of expression for CD11b (official name ITGAM), a marker of myeloid immune cells such as macrophages, granulocytes, and their monocyte progenitors [58]. As compared to controls or *Mafb* KO;*Maf*-heterozygous gonads, there was a dramatic increase in the number of CD11b-bright cells in *Mafb-*heterozygous;*Maf* KO and double KO gonad/mesonephros complexes in both sexes (Figure 5E-H and M-N). These findings suggest that *Maf* is required to suppress supernumerary CD11b-bright immune cells in the gonad and mesonephros.

**Figure 5.**
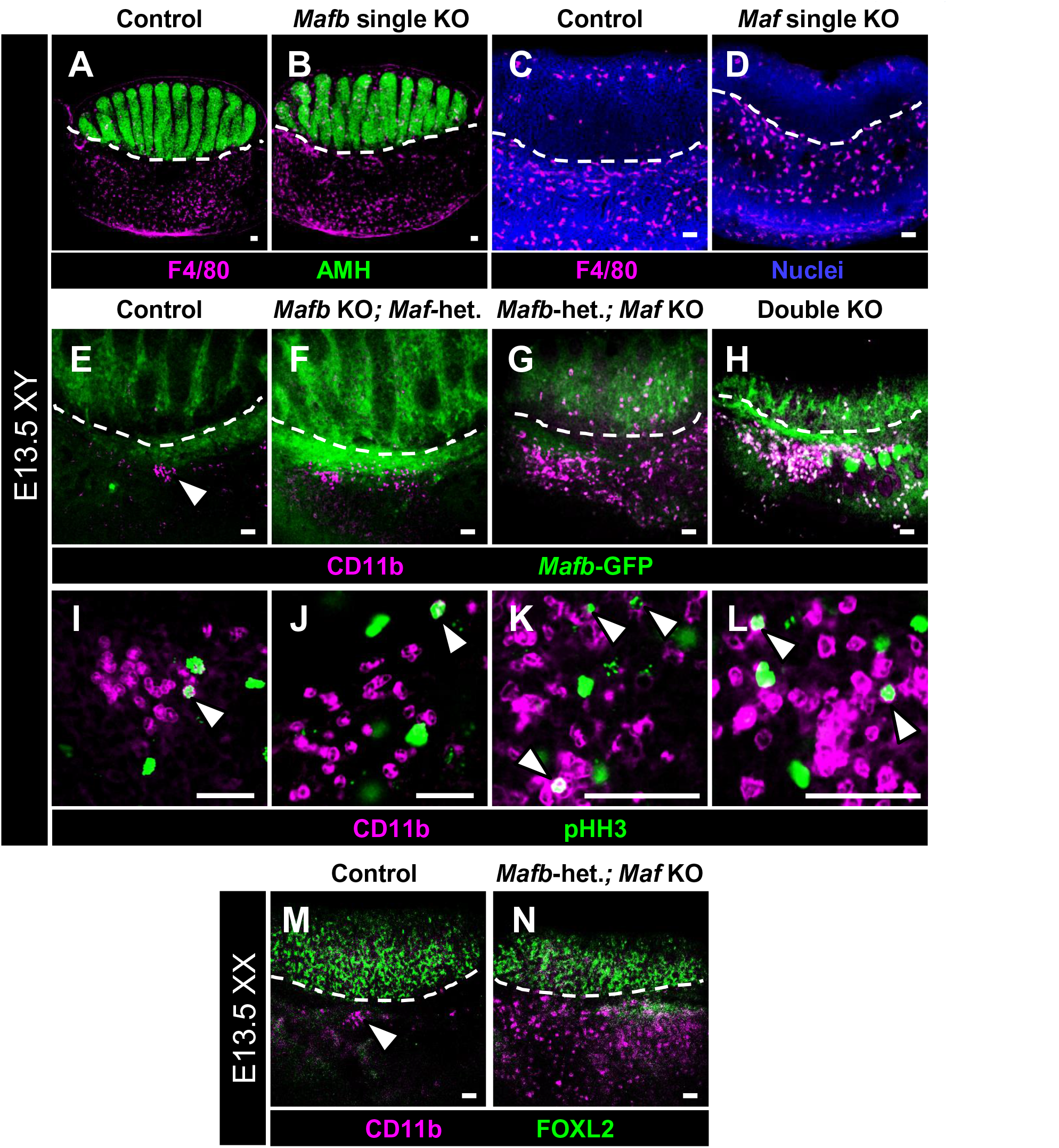
*Maf* mutant gonads display supernumerary CD11b-bright immune cells. Immunofluorescent images of E13.5 XY control (A, C, E, and I), *Mafb* single KO (B), *Maf* single KO (D), *Mafb* KO;*Maf*-heterozygous (F and J), *Mafb-*heterozygous;*Maf* KO (G and K), and double KO (H and L) gonads. Dashed lines mark gonad-mesonephros boundary. (A-D) F4/80+ differentiated macrophages are detected in similar numbers between control (A and C), *Mafb* single KO (B), and *Maf* single KO (D) fetal testes. However, in both *Mafb-*heterozygous;*Maf* KO (G) and double KO (H) testes, there is a dramatic increase in CD11b-bright cells relative to the small clusters of cells (arrowhead in E) found in control samples. (I-L) All genotypes exhibit mitotic (pHH3+) CD11b-bright cells (arrowheads in I-L). (M and N) Compared to E13.5 XX control ovaries (M), which contain a cluster of CD11b-bright cells along the gonad-mesonephros border (arrowhead in M), E13.5 XX *Mafb*-heterozygous;*Maf* KO fetal ovaries (N) possess supernumerary CD11b-bright cells throughout the mesonephros. Scale bars, 50 μm.

To address the source of increased numbers of CD11b-bright cells, we assessed whether these cells were proliferative in the fetal testis, using phosphoHistone H3 (pHH3) to mark mitotic cells. We detected pHH3 expression in CD11b-bright cells in E13.5 control, *Mafb* KO;*Maf*-heterozygous, *Mafb-*heterozygous;*Maf* KO, and double KO testes and mesonephros (Figure 5I-L). These data indicate that differential mitotic activity is unlikely to be the sole source of supernumerary CD11b-bright cells in *Maf* KO embryos.

### Supernumerary immune cells in Maf KO and double KO gonads are monocytes

We next sought to determine the identity of ectopic immune cells in KO gonad/mesonephros complexes. To label and identify all immune cells in the gonad/mesonephros complex, we permanently lineage-traced all hematopoietic-derived cells and their progeny using a *Rosa*-Tomato fluorescent reporter driven by *Vav1*-Cre. At E12.5, when testis-specific vascular remodeling and initial sexual differentiation occur, most Tomato+ cells in the developing gonad-mesonephros complex were F4/80+ macrophages (Figure 6A). However, there was a population of Tomato+ cells that did not express F4/80 (Figure 6A), which had a unique, round morphology as compared to more ramified, dendritic-like macrophages. To confirm these cells were in fact immune cells, we examined expression of CD45, a pan-hematopoietic marker, as well as CD11b and F4/80, and confirmed that gonadal and mesonephric CD11b-bright round cells were immune cells that were not differentiated macrophages (Figure 6B and C).

**Figure 6.**
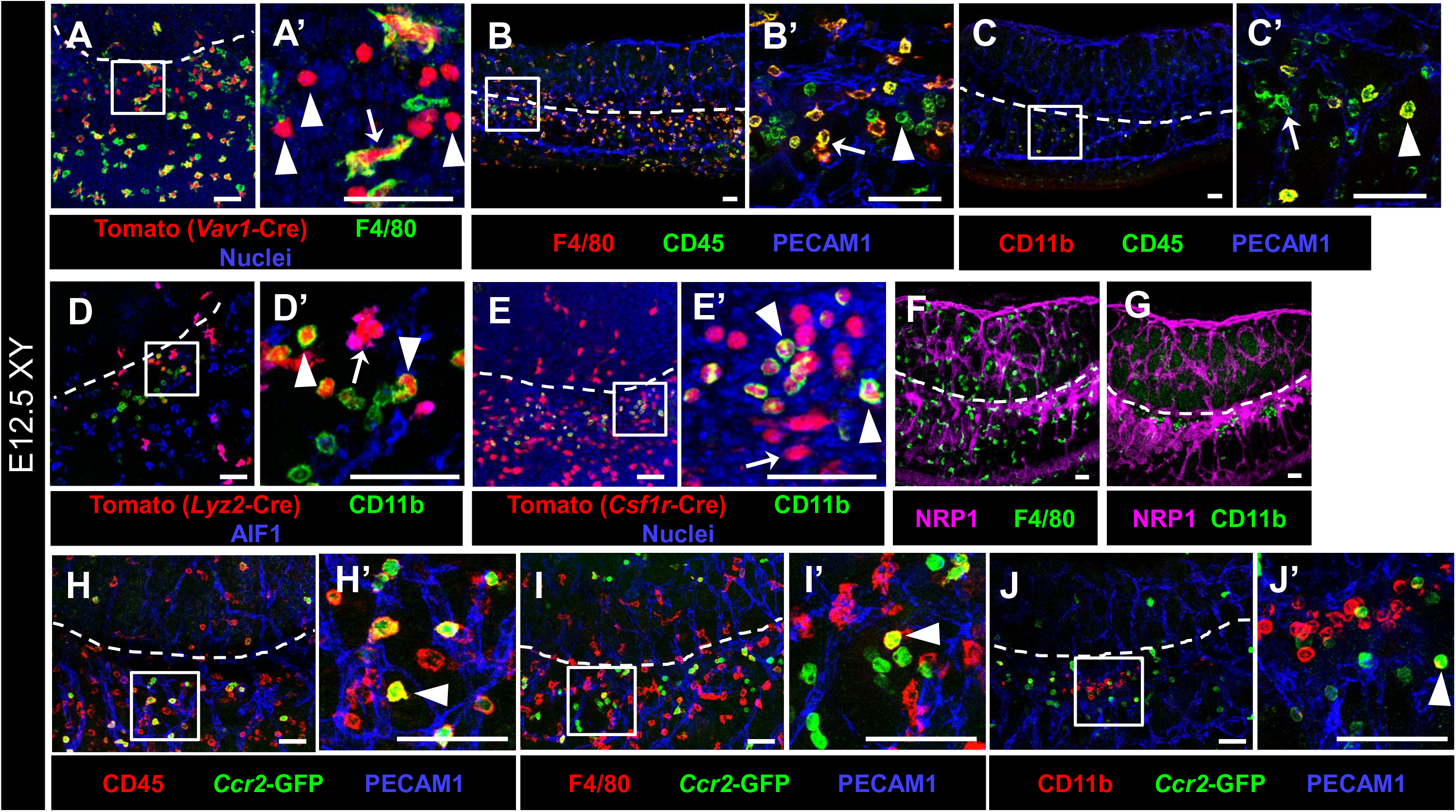
CD11b-bright cells are fetal monocytes specifically localized near the gonad-mesonephric vascular plexus. Immunofluorescent images of E12.5 XY *Vav1*-Cre; *Rosa*-Tomato (A), wild-type C57BL/6J (B,C,F,G), *Lyz2*-Cre; *Rosa*-Tomato (D), *Csf1r*-Cre; *Rosa*-Tomato (E), and *Ccr2*-GFP (H-J) gonads. A’-E’ and H’-J’ are higher-magnification images of the boxed regions in A-E and H-J. White dashed lines indicate gonad-mesonephros border throughout the figure. Labeling all hematopoietic cells with Tomato via *Vav1*-Cre (A) reveals both F4/80+ macrophages (arrow) and F4/80-negative round cells (arrowheads). (B,C) Staining with CD45 reveals F4/80-bright and CD11b-dim macrophages (B’ and C’, arrows), as well as F4/80-dim/negative, CD11b-bright round cells (B’ and C’, arrowheads). (D, E) Targeting myeloid cells with *Lyz2*-Cre (D) and monocyte/macrophages with *Csf1r*-Cre (E) reveals Tomato+ macrophages (D’ and E’, arrows; AIF1+ in D’) and CD11b-bright round cells (D’ and E’, arrowheads). (F, G) Whereas F4/80+ macrophages are associated with vasculature throughout the entire gonad-mesonephros complex (F), CD11b-bright cells are specifically localized near the gonad-mesonephric vascular plexus (G). NRP1 labels endothelial cells. (H-K) The monocyte marker *Ccr2*-GFP reveals GFP+/CD45+ cells near the gonad border (H’, arrowhead), which are occasionally F4/80+ (I’, arrowhead) and CD11b-bright (J’, arrowhead), but are mostly only GFP+. Scale bars, 50 μm.

To determine if CD11b-bright cells had a myeloid identity, we lineage-traced myeloid cells and their progeny using *Lyz2*-Cre. Despite a low labeling efficiency (∼20%) in fetal gonad immune cells, we found AIF1+ (also called IBA1) macrophages labeled with the fluorescent lineage reporter (Figure 6D). In addition to Tomato+ macrophages, we also identified Tomato+, AIF1-negative round cells that strongly expressed CD11b (Figure 6D). These data indicate that this population of immune cells in the fetal gonad/mesonephros complex originated from a myeloid lineage.

To investigate further the immune identity of CD11b-bright cells within the myeloid lineage, we utilized *Csf1r*-Cre to lineage-trace monocyte/macrophage cells. We found that CD11b-bright cells were effectively labeled with Tomato (Figure 6E), similar to gonadal and mesonephric macrophages, indicating that CD11b-bright cells were likely monocytes or monocyte-derived cells. Consistent with a non-macrophage identity, this subpopulation of hematopoietic-derived cells did not express colony stimulating factor receptor 1 (CSF1R), AIF1, neuropilin 1 (NRP1), MRC1 (also called CD206) or *Cx3cr1*-GFP (Supplemental Figure S5G-K), markers we previously reported as markers of fetal testicular macrophages [8].

In contrast to tissue-resident F4/80+ macrophages, which were found throughout the gonad and mesonephros, *Mafb-*GFP-positive, CD11b-bright cells were specifically localized near vasculature at the vascular plexus of the gonad-mesonephros border in both sexes, as well as the developing coelomic artery in males (Figure 6F and G; Supplemental Figure S5G and H). In addition, round CD11b-bright cells possessed a characteristic horseshoe-shaped nuclear morphology (Supplemental Figure S5L), indicating that these cells were monocytes. To address this hypothesis, we examined a marker of monocytes, *Ccr2-*GFP, and found numerous round GFP+ cells in the gonad, which were mostly localized near the vascularized gonad-mesonephros border at E12.5 (Figure 6H-J). GFP+ cells, as expected, expressed CD45 (Figure 6H), and did not express F4/80 except in a small number of cells (Figure 6I). However, unexpectedly, GFP+ cells did not yet express CD11b, possibly suggesting that these cells are myeloid progenitor cells. Overall, our data suggest that the round ectopic cells observed in *Maf* KO and double KO gonad/mesonephros complexes are likely monocytes or, alternatively, a myeloid progenitor cell.

### MAFB and MAF are expressed in macrophages but not in monocytes

To address the potential mechanisms of how *Mafb* and *Maf* regulate hematopoiesis, we examined the expression of MAFB and MAF within gonadal and mesonephric immune cells. We found that both MAFB and MAF were expressed in F4/80+ macrophages in the gonad/mesonephros complex, which were likely of yolk sac origin, as early as E10.5 (Supplemental Figure S6A and B). At E12.5, during testicular morphogenesis, MAFB and MAF were both expressed in F4/80-positive macrophages (Supplemental Figure S6C and D), but neither MAFB nor MAF were expressed in CD11b-bright monocytes in the fetal gonad and mesonephros (Supplemental Figure S6E and F). The co-expression of MAFB and MAF in F4/80+ cells was maintained at E13.5 (Supplemental Figure S6G).

### Mafb and Maf mutation disrupts Leydig cell differentiation and immune gene expression

We next sought to determine if other interstitial cell types, apart from vasculature, were disrupted by *Mafb* and *Maf* loss of function. To assess specifically the effects of *Maf* loss of function on interstitial cells, we FACS-purified *Mafb-*GFP-positive cells, which include interstitial mesenchymal and immune cells [9, 54], from E12.5 *Mafb-*heterozygous;*Maf* KO versus control fetal testis-mesonephros complexes and performed microarray transcriptomic analyses (Table 1; Supplemental Table S1). We found among the top 20 downregulated genes were all the major components of the Leydig cell steroidogenic pathway: *Star*, *Cyp11a1*, *Hsd3b1*, and *Cyp17a1* were all reduced in *Maf*-mutant interstitial cells (Figure 7A; Table 1). Additionally, other Leydig-specific genes such as *Insl3* and *Ren1* were downregulated (Figure 7A; Table 1), suggesting there was a reduction in Leydig cell number as opposed to a specific disruption of the steroidogenic pathway within a normal number of Leydig cells. In immunofluorescence analyses, control testes contained Leydig cells throughout the interstitium (Figure 7B), but there was a reduction in Leydig cell number in *Mafb-*heterozygous;*Maf* KO testes (Figure 7C). We found that double KO gonads also exhibited a reduced Leydig cell number relative to controls in immunofluorescence assays (Figure 7D and E).

**Figure 7.**
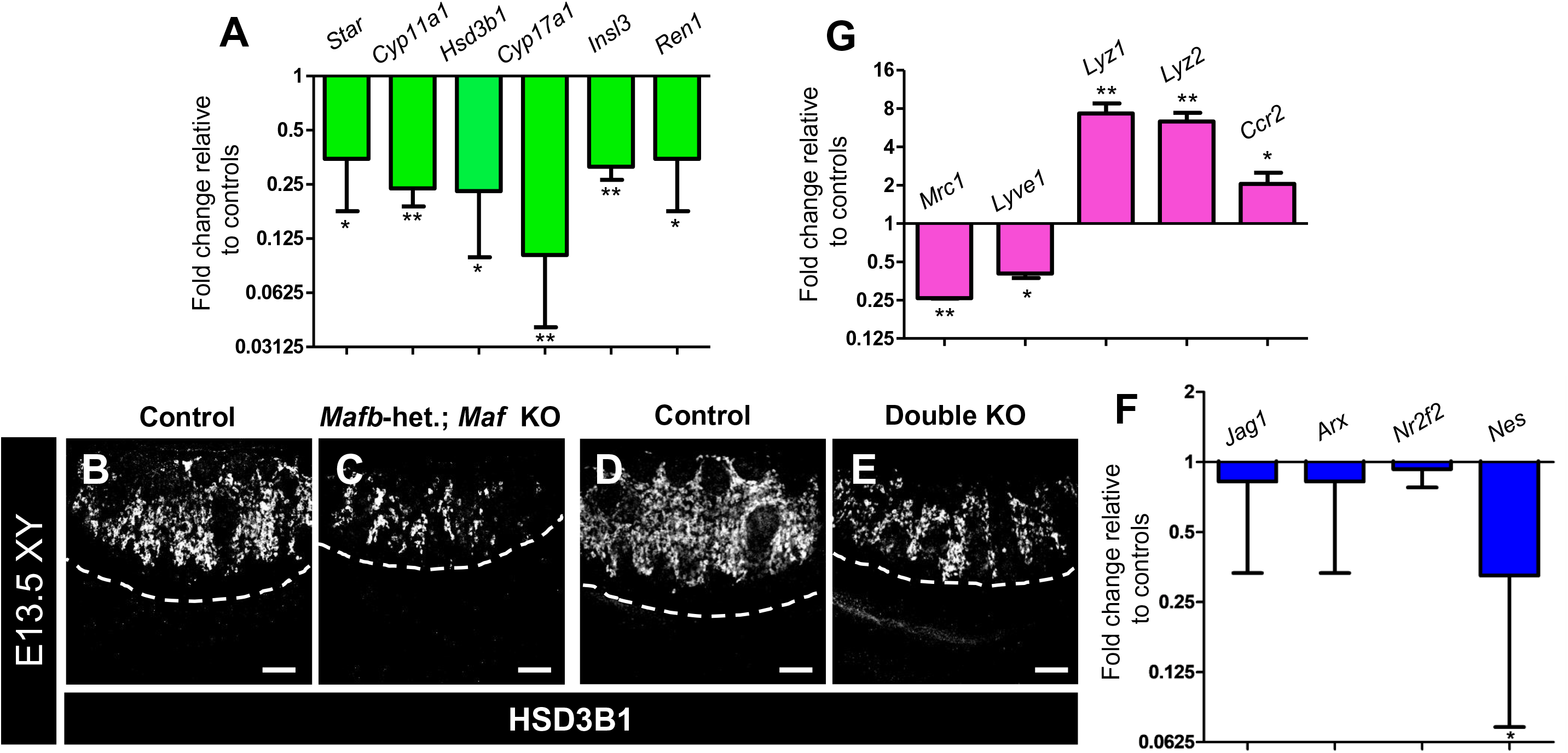
*Maf* loss of function causes disruptions in Leydig and immune cell differentiation. (A) Graph showing gene expression fold change from microarray gene expression analysis of *Mafb-*heterozygous;*Maf* KO GFP+ interstitial cells FACS-purified from E12.5 XY gonad-mesonephros complexes, showing reduction in Leydig cell gene expression. (B-E) Immunofluorescent images of E13.5 XY control (B,D), *Mafb*-heterozygous;*Maf* KO (C), and double KO (E) gonads, showing reduction of HSD3B1+ Leydig cells in KO gonads. White dashed lines indicate gonad-mesonephros border. Scale bars, 100 μm. (F) qRT-PCR analyses for interstitial progenitor-specific gene expression in E13.5 XY *Mafb*-heterozygous;*Maf* KO gonads relative to controls. (G) Microarray analysis of gene expression change in *Mafb-*heterozygous;*Maf* KO interstitial cells (*Mafb-*GFP+) FACS-purified from E12.5 XY gonad-mesonephros complexes (same dataset as in A), showing reduction in M2 macrophage gene expression and increase in genes associated with degradative myeloid cells and monocytes. All graph data are represented as mean +/- SD. *, *P*<0.05; **, *P*<0.01 (Student t-test).

**Table 1.**
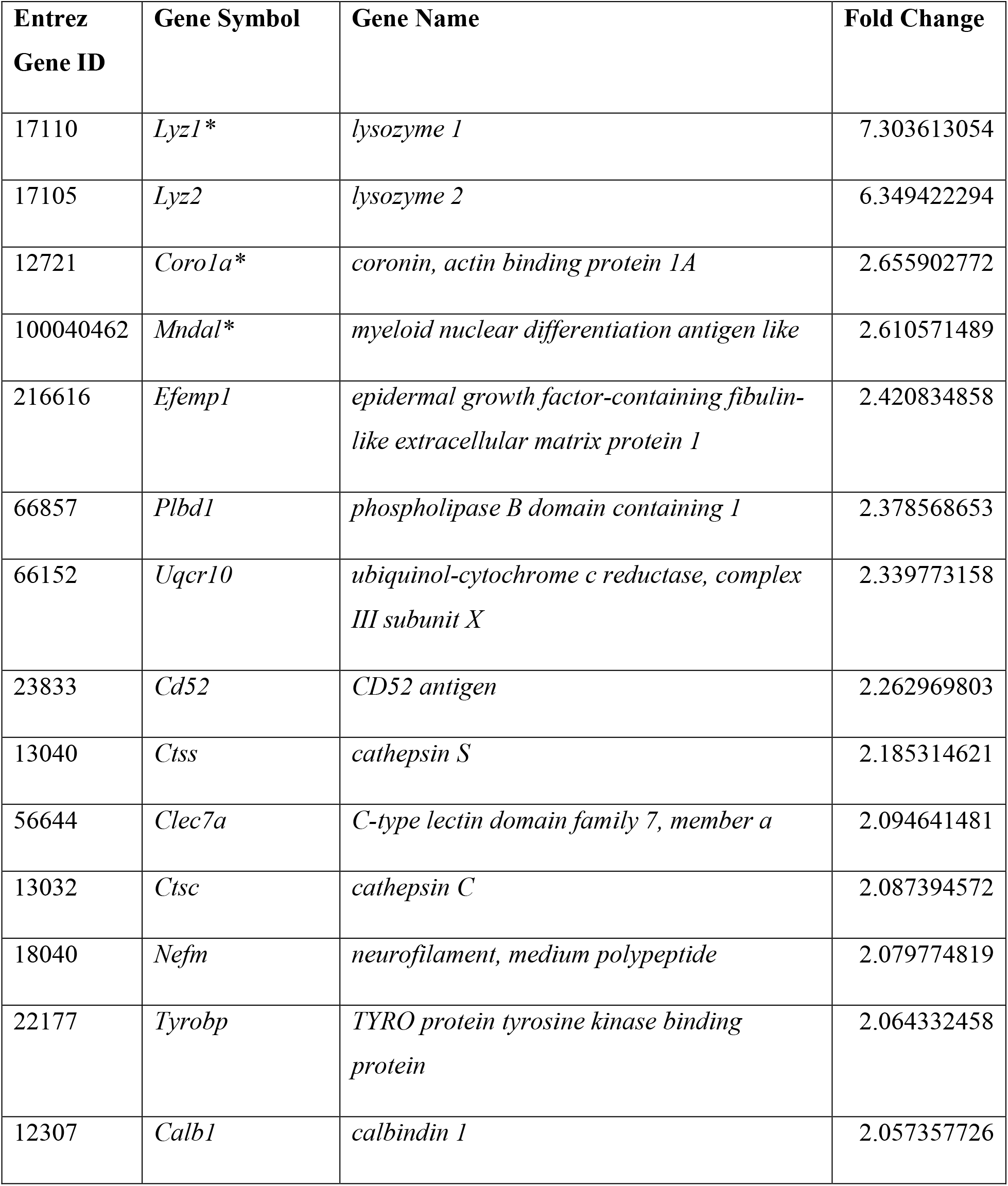
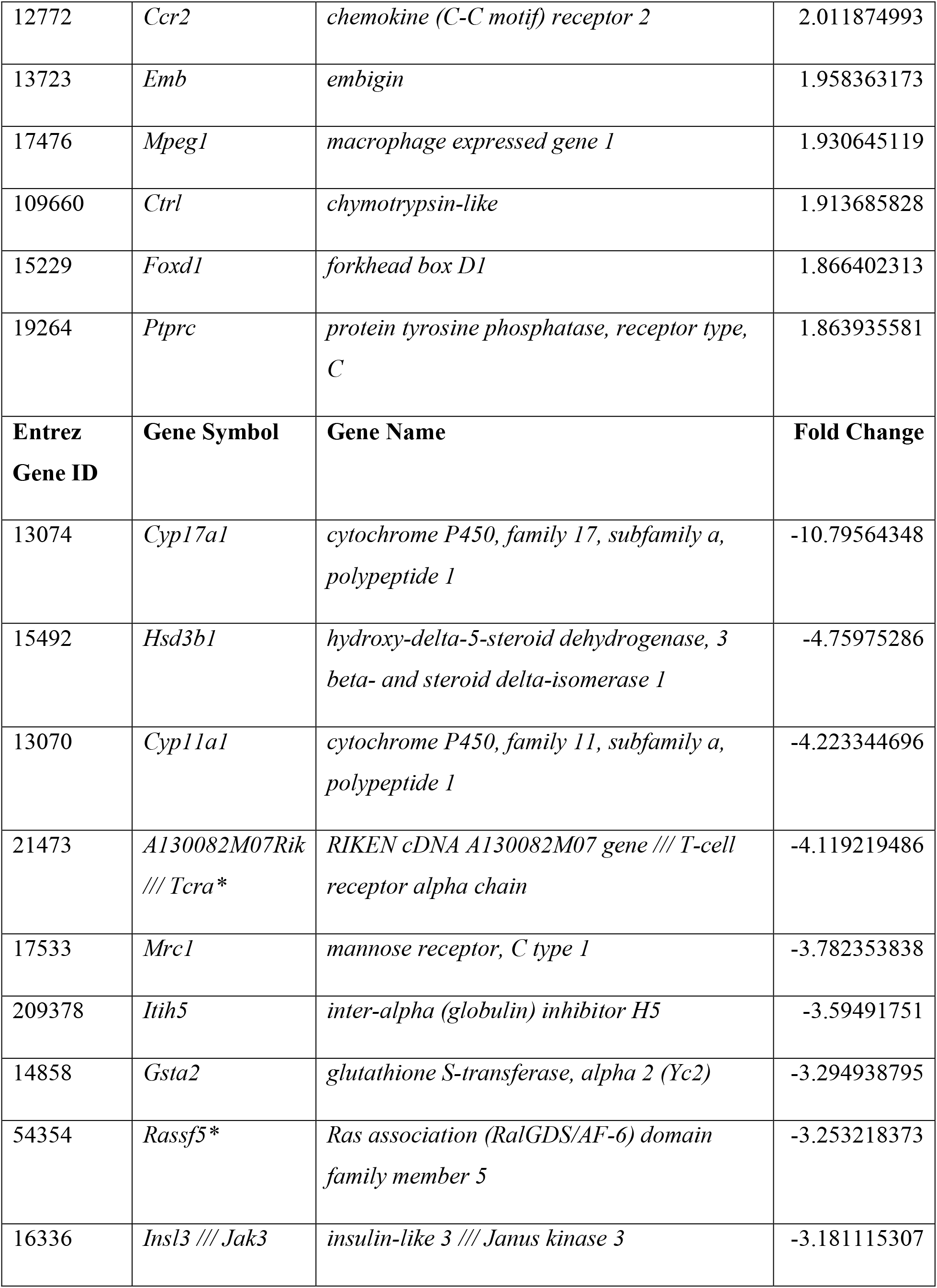
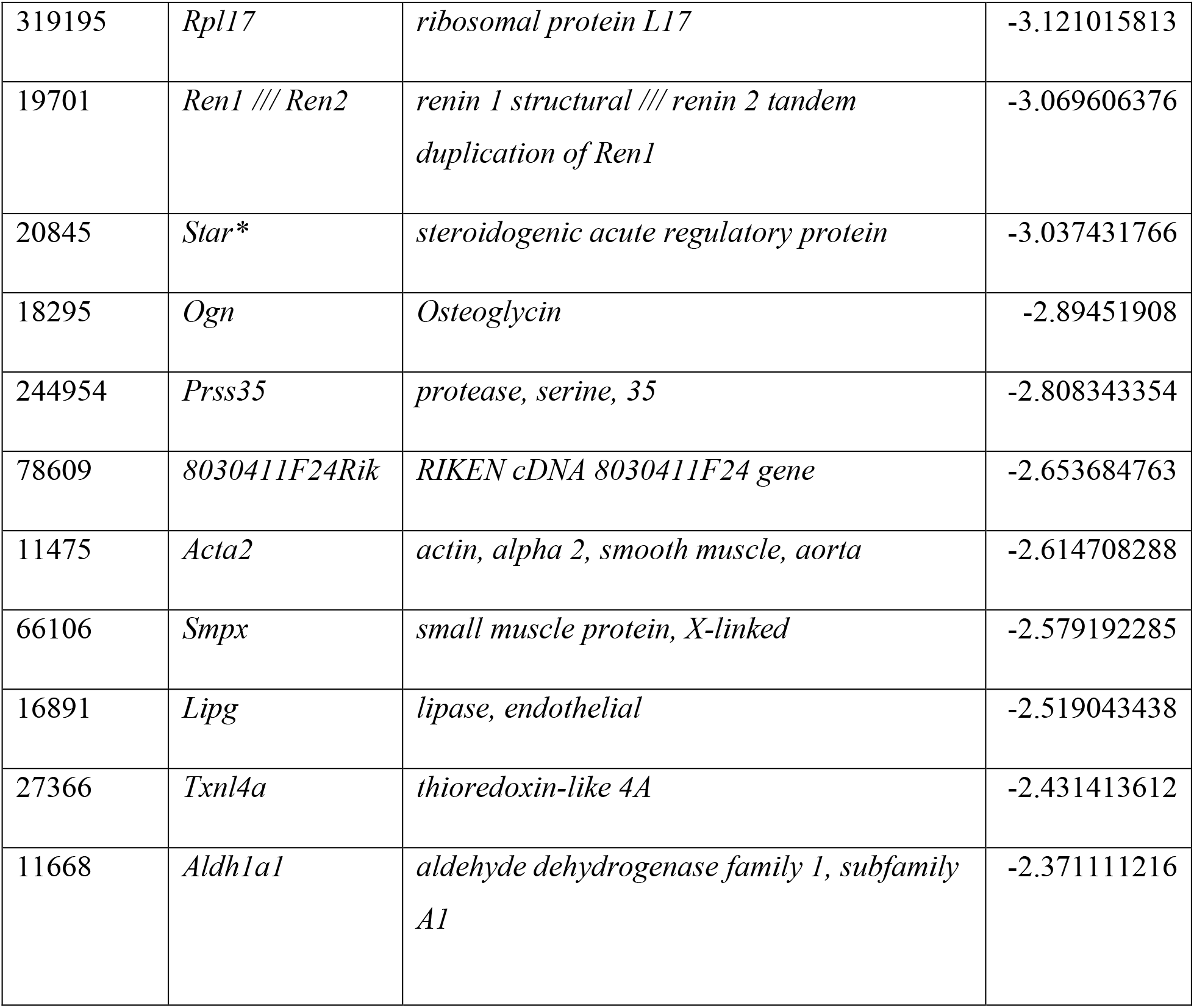
Upregulated and downregulated genes in *Mafb*-heterozygous;*Maf* KO interstitial (*Mafb*-GFP-expressing) cells. Table shows the top 20 upregulated and top 20 downregulated genes in FACS-purified GFP+ cells from E12.5 XY *Mafb*-heterozygous;*Maf* KO (*Mafb^GFP/+^*;*Maf^-/-^*)versus control (*Mafb^GFP/+^*;*Maf^+/-^*) testis and mesonephric tissue. Asterisks denote genes that were represented by two probe sets within the top 20 upregulated or downregulated probe sets; the most upregulated or downregulated probe set for that gene is listed.

To determine any effects on Leydig progenitors, we performed qRT-PCR analyses on E13.5 XY control and *Mafb-*heterozygous;*Maf* KO gonads for several interstitial progenitor-specific genes, such as *Jag1*, *Arx*, *Nr2f2* (also called *COUP-TFII*), and *Nes* (*Nestin*). We only found a reduction in *Nes* expression (Figure 7F), which is specific to a subset of perivascular progenitor cells [10], indicating there may be some defects in vascular-mesenchymal interactions or Leydig cell differentiation, but there do not appear to be any widespread, general defects in the establishment of progenitor populations in KO fetal testes.

As opposed to downregulated genes, which were mostly associated with Leydig cells, most upregulated genes in *Mafb-*heterozygous;*Maf* KO cells were associated with macrophage and monocyte immune function. While genes normally expressed in M2-type tissue-resident macrophages, such as *Mrc1* (CD206) and *Lyve1* were significantly downregulated, genes associated with monocytes, such as *Ccr2* and *Ptprc* (CD45), and degradative activity of myeloid cells, such as *Lyz1*, *Lyz2*, and cathepsin-encoding genes *Ctss* and *Ctsc,* were upregulated (Figure 7G; Table 1; Supplemental Table S1). In addition, the gene encoding the actin regulatory protein Coronin1a (*Coro1a*), involved in forming the phagolysosome, was also significantly upregulated in *Mafb-*heterozygous;*Maf* KO cells, along with other less well-characterized genes associated with myeloid or immune function (Supplemental Table S1), suggesting that the ectopic immune cells in *Maf* KO gonads were phagocytic and had degradative activity.

### Leydig cell reduction in KO gonads is likely due to hypervascularization

Our previous work demonstrated that Leydig progenitors in the fetal testis were maintained by vasculature, and the number of differentiated Leydig cells was increased when vascularization was inhibited ex vivo [10]. The converse situation occurred in *Maf* single KO and double KO testes, in which there was hypervascularization; therefore, we hypothesized that the reduction of fetal Leydig cells in mutant gonads was potentially a secondary effect in which excess or dysregulated vascularization inhibited the ability of interstitial cells to differentiate into Leydig cells. To address this hypothesis, we manipulated vascularization in XY gonads via the addition of PDGF-BB in ex vivo culture. PDGF signaling regulates vascularization in the fetal testis, and addition of PDGF-BB to both XX and XY gonads ex vivo induced vasculature [7, 59]. After addition of PDGF-BB, we found that vascular patterning was disrupted, as evident by increased blood vessels in the coelomic surface of the gonad (Figure 8A and B). Along with increased surface vasculature, we found that there was a reduction in Leydig cells in PDGF-BB-treated fetal testes (Figure 8B). Using qRT-PCR, we found that the expression of the endothelial marker *Cdh5* was not increased, indicating that the overall number of endothelial cells was likely not changed, but that instead vascular remodeling and patterning were affected. Additionally, the expression of Sertoli cell (*Sox9*, *Amh*) and germ cell (*Ddx4*) genes was not significantly different in PDGF-BB-treated testes (Figure 8C). Consistent with immunofluorescence data for CYP11A1, expression of *Cyp11a1*, *Hsd3b1*, and *Cyp17a1* mRNA were all significantly reduced in PDGF-BB-treated testes relative to controls (Figure 8C), indicating a reduction in Leydig cells. Similar to PDGF-BB treatment, we also saw that Leydig cell number was reduced in another scenario in which vascular patterning was experimentally disrupted, such as culturing in 10% FBS instead of 5% FBS (Figure 8D-F) and in the presence of VEGFA (Supplemental Figure S7). Therefore, our data suggest that dysregulated vasculature in *Maf* KO or double KO testes is the likely cause of reduced Leydig cell number in mutant gonads.

**Figure 8.**
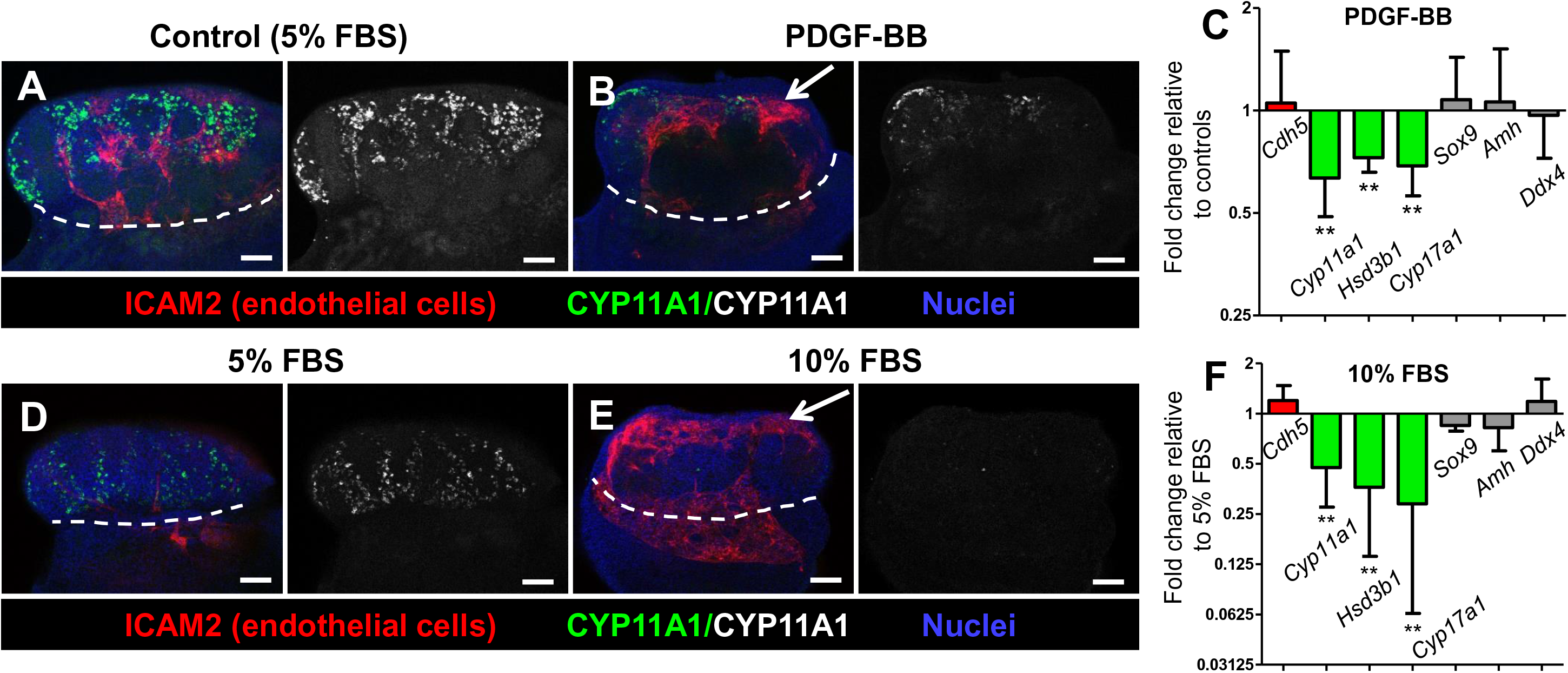
Disrupted vascular patterning in the fetal testis is associated with reduced Leydig cell differentiation. Immunofluorescent (A,B,D,E) and qRT-PCR (C,F) analyses of 48-hour ex vivo gonad culture of E12.5 CD-1 gonads, showing that disruptions in vascular patterning (arrows in B and E) caused by either PDGF-BB treatment (A-C) or increase in FBS concentration in the culture media (D-F) resulted in a decreased number of Leydig cells without effects on Sertoli or germ cells. White dashed lines indicate gonad-mesonephros border. Scale bars, 100 μm. All graph data are represented as mean +/- SD. **, *P*<0.01 (Student t-test).

To address whether factors other than disrupted vasculature are the cause of reduced Leydig cells in PDGF-BB-treated testes, we performed PDGF-BB gonad culture experiments in the presence of the vascular inhibitor VEGFR-TKI II to eliminate vasculature during PDGF-BB treatment. We found that there was no significant difference in Leydig cell gene expression in PDGFBB+VEGFR-TKI-II-treated versus VEGFR-TKI-II-alone treated testes (Supplemental Figure S8A-E), indicating that vasculature, or the lack of vasculature in this case, was the main driver of the Leydig cell phenotype in PDGF-BB culture experiments.

Finally, to address whether there is any potential miscommunication between Sertoli and Leydig cells that could cause Leydig cell dysregulation in KO gonads, we examined several pathways that involve Sertoli-Leydig crosstalk. qRT-PCR analyses for the Pdgf pathway (*Pdgfa* and *Pdgfra*) and Notch pathway (Notch interstitial target genes *Hes1*, *Hey1*, and *Heyl*) did not reveal any defects in *Mafb*-heterozygous;*Maf* KO gonads (Supplementary Figure S8F). We also examined the desert hedgehog (DHH) pathway; while we did not see any effect on mRNA levels of *Dhh*, which is expressed by Sertoli cells [60, 61], in *Mafb*-heterozygous;*Maf* KO gonads, we did observe a significant reduction in the expression of *Ptch1* (*patched 1*), which encodes the DHH receptor and is expressed in the fetal testis interstitial compartment [61] (Supplementary Figure S8F).

## Discussion

In this study we have uncovered new roles for the transcription factors MAFB and MAF (C-MAF) in gonadal development and hematopoiesis. Our data demonstrate that *Mafb* and *Maf*, acting redundantly, regulate immune cell fate and vascular remodeling that are required for testicular differentiation and morphogenesis. In double KO gonads, we observed a significant increase in the number of monocytes, which was associated with multiple perturbations in gonadal development, including testicular hypervascularization, testis cord abnormalities, Leydig cell deficits, and a reduced number of germ cells in both sexes. While mutations in the *Drosophila* large Maf gene *traffic jam* caused gonad morphogenesis defects via disruption of cell adhesion molecules [38], here we found no evidence from transcriptome data that this was the case in mice. Instead, our results suggest that aberrant gonad development in mice was caused by *Maf*-dependent changes in hematopoiesis that resulted in disruption of vascular remodeling. These results support a broadly emerging idea that vasculature and the balance of immune cell types are critical for mammalian organogenesis.

The family of large Maf transcription factors has been described, in multiple contexts, as critical regulators of cellular differentiation during organogenesis [62, 63]. In hematopoiesis, both *MafB* and *Maf* have significant roles in the fate of myeloid cells. MAFB directly interacts with the DNA-binding domain of ETS-1, thereby repressing erythroid differentiation in pluripotent myeloid cells [64]. Through transduction of *Mafb* in hematopoietic precursor cells, *Mafb* has been further shown to promote formation of myeloid colonies and macrophage differentiation [65, 66]. *Maf* also possesses a regulatory role in myelomonocytic differentiation, though its involvement is currently not defined as definitively as *Mafb*’s monocytic promotion. Induced expression of *Maf* resulted in the accumulation of monocytes and macrophages, followed by their eventual apoptosis [67]. More recently, analysis of *Maf*-deficient embryos revealed that they are anemic due to deficiencies in macrophage functions essential for maintenance of erythroblastic island formation and functional erythrypoiesis [28]. Indeed, *Maf* has been repeatedly observed to regulate expression of various genes encoding immune cytokines, such as *Il4* and *Il21* [68, 69].

Our data, in conjunction with previous studies, point to a scenario in which lack of *Maf*, or lack of both *Maf* and *Mafb*, results in a bias in hematopoietic differentiation toward a gonadal monocyte fate. In our study, the F4/80-positive macrophage population was comparable between control, *Mafb* single KO, and *Maf* single KO gonads, indicating that individual Maf genes are not required for tissue-resident macrophage differentiation in the gonad. However, the CD11b-bright population of monocytes was dramatically increased in *Mafb*-heterozygous;*Maf* KO and double KO gonads. A previous study demonstrated that mutation in *Mafb* and *Maf* disassociated cell cycle activity from differentiation in hematopoietic cells, resulting in extensive proliferation of mature monocytes and macrophages [27], which rarely occurs in normal development. In addition, *Maf* also is involved in inducing apoptosis of CD11b-expressing monocytic and myeloid cells [67]. Given the well-characterized roles of Maf factors in cell fate determination, we propose that the Maf family of genes normally suppresses the differentiation or survival of CD11b-positive monocytes from a hematopoietic progenitor population. This idea is consistent with our observation that CD11b-positive monocytes are MAFB-negative and MAF-negative. An in-depth analysis of myeloid cell populations in *Maf* KO and double KO gonads could uncover further roles for this gene in regulating cell fate decisions during organogenesis and organ homeostasis.

Blood vessels form an intricate and interconnected network that is critical for sustaining functional organs via oxygen and nutrient supply to tissues. Prior to vascular function in delivering blood flow, embryonic endothelial cells (ECs) and nascent vessels play a general role in promoting organogenesis, as has been reported in liver, testis, and pancreas [7, 10, 70–72]. ECs are important components of essential niches for stem cell self-renewal versus differentiation during organogenesis [70], such as during pancreas development, in which pancreatic progenitors rely on EC-supplied EGFL7 for renewal and maintenance [73]. Our previous results showed that EC-derived Notch signaling is essential for maintaining fetal Leydig progenitors in mice, whereby both vascular inhibition and inactivation of Notch signaling induced excess fetal Leydig cell differentiation and loss of Nestin-positive interstitial progenitor cells [10]. Conversely, stimulation of Notch signaling by zearalenone administration *in utero* (likely mediated via the vascular- and perivascular-associated Notch receptors NOTCH1 and NOTCH3) inhibited differentiation of fetal Leydig cells in rats [74]. Therefore, aberrant vascularization in double KO gonads likely disrupted vascular-mesenchymal interactions responsible for promoting differentiation of interstitial cells and establishing a niche for Leydig cell progenitors. This paradigm applies to both double KO gonads and *Maf-*intact gonads in which we experimentally disrupted testicular vascularization ex vivo, demonstrating the importance of proper vascular remodeling on testicular organogenesis. Our results here do not point toward disruption of Notch as a potential mechanism in KO gonads, as interstitial Notch target gene expression was unaffected. However, we did observe a reduction in *Nes* expression, which is expressed in perivascular progenitor cells, indicating that there are some underlying defects in vascular interactions. We also observed a reduction in *Ptch1* expression, which encodes the receptor for the Hedgehog ligand DHH that is essential for fetal Leydig cell differentiation, implicating a disruption in Hedgehog signaling as a potential cause of Leydig cell defects in KO gonads. We will add the caveat that our microarray transcriptome analyses were performed at E12.5, which may be too early to detect some Leydig-cell-specific genes reliably at that stage [75]; therefore, it should be kept in mind that some of our transcriptome findings may be based on differences in low-level expression.

There were clear differences in the phenotype here, in which gonadal myeloid cells were dramatically increased in number due to *Maf* mutation, versus our previous report in which we ablated myeloid cells via a Cre-mediated approach [8]. Specifically, a major difference was observed regarding the vascular network of the mesonephric vascular plexus, from which testicular vasculature (especially the coelomic artery) is derived due to migration of freed mesonephric endothelial cells into the gonad [55, 57]. Here we found that the existing vascular network of the mesonephric vascular plexus was excessively degraded, leading to a dramatic disruption in vascular patterning and hypervascularization of the testis. In contrast, depletion of myeloid cells in our previous study resulted in a poorly remodeled vascular plexus, in which a reduced number of migrating endothelial cells failed to generate a coelomic artery. However, in both cases testicular organ architecture and vascularization were disrupted, leading to aberrant cord formation and, in this study, a disruption of Leydig cell differentiation. Overall, these results demonstrate that a proper balance of immune cell number is critical for regulating vascular remodeling that establishes the morphogenetic and differentiation programs of the developing testis.

Our transcriptome data indicated that genes encoding degradative enzymes, such as lysozymes and cathepsins, were upregulated in *Mafb*-heterozygous, *Maf* KO gonads, and ectopic monocytes were specifically localized near vasculature in the mesonephros region near the gonad border. Therefore, it is likely that supernumerary monocytes in KO gonads led to a disruption of vascular remodeling by excessive, dysregulated breakdown of vasculature in areas such as in the gonad-mesonephros vascular plexus. A growing body of work has shown that monocyte-macrophage cells are critical to support proper vascular remodeling and growth in development [76], and here we show that extensive hypervascularization occurs in *Maf* KO and double KO gonads that possess supernumerary monocytes. An increased accumulation of CD11b+ myeloid cells in this study and its association with hypervascularization is reminiscent of tumor models, in which myeloid cell recruitment is linked to tumor vasculature and growth recovery after radiation [77]. It is well-accepted that CD11b+ myeloid cells have proangiogenic activity to promote the formation of tumor vasculature [78], but here we propose that monocytes can also drive disruptions in vascular remodeling when dysregulated in developing organs. One possibility for monocyte action in the gonad is CD11b–mediated binding of monocytes to endothelial ICAM1, which contributes to vascular sprouting in liver sinusoids and portal space after partial hepatectomy [79, 80]. Therefore, we posit that the dramatic, dysregulated increase of CD11b-positive monocytes in double KO gonads leads to hypervascularization resulting from a disruption in the balance of vascular remodeling versus breakdown during testis differentiation.

Therefore, our data add to evidence supporting the idea that ECs are essential regulators of organ formation, which when disrupted, can lead to developmental abnormalities. In humans, birth defects of various organ structures, such as in the gut and limbs, have been linked with vascular disruption [81], indicating that further investigation into the role of blood vessels during fetal development will be an important area of future research. Our findings here also highlight the non-immune roles that vascular and hematopoietic/immune cells play during mammalian gonad development. Studies from other model systems suggest that non-immune, developmental functions for immune cells are evolutionarily conserved across the animal kingdom. In *Xenopus* embryos, targeted ablation of macrophages resulted in developmental defects such as disrupted limb morphogenesis and early death [82]. In *Drosophila*, the macrophage-like hemocyte lineage plays important roles in organogenesis, such as in central nervous system morphogenesis [83, 84], where it acts through modulation of extracellular proteins and clearance of apoptotic cells, and in the intestinal stem cell niche, where hemocytes regulate stem cell proliferation via BMP signaling [85]. In the study of *Drosophila traffic jam* mutants [38], it was not addressed whether there was a change in hemocytes; therefore, it is unclear whether immune cells played a role in the *traffic jam* gonad phenotype. Interestingly, limb regeneration in adult salamanders and fin, heart, and axonal regeneration in zebrafish all require myeloid cells [86–89], and perhaps also in heart repair in mouse injury models [90]. Regeneration studies in diverse species have led to an emerging idea that a proper immune response is essential for both organ formation and regeneration [91]. Increasing evidence supports the idea that myeloid cells, such as monocytes and macrophages, play broad and evolutionarily conserved roles in organogenesis via their extensive repertoire of cellular and molecular functions, one of which is regulating vascular and tissue formation and function. Further investigation into the links between immune cell activity, vascularization, and morphogenesis will be critical for a deeper understanding of organogenesis and fetal development.

## Acknowledgments

We thank S. Takahashi, L. Goodrich, I.C. Ho, H.L. Grimes, and R. Lang for mice; we also thank K. Morohashi and D. Wilhelm for antibodies.

## Conflict of interest

None declared.

## Author Contributions

SL conducted experiments, performed data analyses, co-wrote the original manuscript, and edited the manuscript. XG and AH conducted experiments, co-wrote the original manuscript, and edited the manuscript. EH conducted experiments and edited the manuscript. BC supervised the project, acquired funding, and edited the manuscript. TD supervised the project, acquired funding, performed experiments, co-wrote the original manuscript, and edited the manuscript.

## Data availability

Raw data associated with microarray transcriptome analyses are publicly available at the Gene Expression Omnibus (GEO) under accession number GSE41715. Other data underlying this article will be shared on reasonable request to the corresponding author.

## Notes

ǂ This work was supported by the National Institutes of Health (R37HD039963 to BC, R35GM119458 to TD, R01HD094698 to TD, F32HD058433 to TD); March of Dimes (1-FY10-355 to BC, Basil O’Connor Starter Scholar Award 5-FY14-32 to TD); Lalor Foundation (postdoctoral fellowship to SL); and Cincinnati Children’s Hospital Medical Center (Research Innovation and Pilot funding, Trustee Award, and developmental funds to TD).

### Competing Interest Statement

The authors have declared no competing interest.

