## Supplemental Data for "Loss of *Mafb* and *Maf* distorts myeloid cell ratios and disrupts fetal mouse testis vascularization and organogenesis"

### Supplementary Material

#### Supplementary Data Legends

**Supplemental Figure S1. Male gonadal sex differentiation occurs normally in the absence of *Mafb* or *Maf*.** Immunofluorescent images of E11.5 (A-C), E12.5 (D-F), and E13.5 (G-O) XY control (A,D,G,J,M), *Mafb* single KO (B,E,H,K,N), and *Maf* single KO (C,F,I,L,O) gonads. White dashed lines indicate gonad-mesonephros border throughout all Supplemental Figures. PECAM1 labels germ cells and endothelial cells. (A-C) As in control (A) E11.5 fetal testes, *Mafb* single KO (B) and *Maf* single KO (C) fetal testes contain SOX9+ Sertoli cells. (D-F) At E12.5, control testes (D) had testis cords containing NR5A1+ (also called SF1) Sertoli cells and PECAM1+ germ cells. In *Mafb* single KO (E) and *Maf* single KO (F) mutants, testis cords were irregularly shaped, variably depleted of germ cells (arrowheads in E,F), or were abnormally small (arrows in F). Solid white lines denote testis cords in D-F. As compared to control (G,J,M) E13.5 fetal testes, *Mafb* single KO (H,K,N) and *Maf* single KO (I,L,O) fetal testes showed recovery of normal testis cord structure (AMH labels Sertoli cells) and germ cell (DDX4+) numbers; contained normal numbers of CYP17A1+ Leydig cells; and underwent testis-specific vascularization. However, *Maf* single KO fetal testes (O) displayed a hypervascularized coelomic surface (arrowhead in O), discontinuous mesonephric vascular plexus (arrow in O), and occasional fused testis cords (bracket in O). Scale bars, 100 μm.

**Supplemental Figure S2. Female gonadal sex differentiation and initial ovarian differentiation occur normally in the absence of *Mafb* or *Maf*.** Immunofluorescent images of E13.5 (A-C) and E14.5 (D-F) XX control (A,D), *Mafb* single KO (B,E), and *Maf* single KO (C,F) gonads. Similar to control XX (A,D) fetal ovaries, *Mafb* single KO (B,E) and *Maf* single

KO (C,F) fetal ovaries possess FOXL2+ granulosa cells and SYCP3+ meiotic germ cells throughout the gonad. Note that there are fewer SYCP3+ cells in *Maf* single KO ovaries. Scale bars, 100 μm.

**Supplemental Figure S3. Germ cell defects in KO gonads are not caused by defects in proliferation, cell cycle status, apoptosis, or sex determination.** (A-H) Immunofluorescence images of E12.5 XY (A-D) and E13.5 XX (E-H) control (A,C,E,G) and double KO (B,D,F,H) gonads, showing comparable levels of mitosis (phosphoHistone H3; pHH3+), apoptosis (cleaved CASP3+), and cell cycle status (MKI67+) in germ and somatic cells of control and double KO gonads. Scale bars, 100 μm. (I and J) qRT-PCR analyses of E11.5 XY (I) and E13.5 XX and XY (J) *Mafb*-heterozygous;*Maf* KO versus control gonads, showing comparable expression of factors involved in germ cell migration (*Cxcl12* and *Kitl*) and sex determination (*Sox9*, *Foxl2*, and *Wnt4*). Graph data are represented as mean +/- SD.

**Supplemental Figure S4. Busulfan-mediated depletion of germ cells does not disrupt testis morphogenesis at E13.5.** Immunofluorescent images of E13.5 fetal testes exposed at E10.5 to vehicle control (A and C) or busulfan (B and D). Control testes (A and C) containing germ cells displayed normal testis cord structure and vasculature. Although busulfan-treated testes (B and D) exhibited a drastic depletion of CDH1+ or PECAM1+ germ cells, they exhibited normal vascularization and testis cord architecture. Scale bars, 100 μm.

**Supplemental Figure S5. Supernumerary *Mafb*-GFP-positive CD11b-bright immune cells in *Maf*-mutant fetal gonads and mesonephroi are likely monocytes.** (A-F) Immunofluorescent images of E13.5 XY (A-D) control (A), *Mafb* KO;*Maf*-heterozygous (B), *Mafb*-heterozygous;*Maf* KO (C), double KO (D), XX (E and F) control (E), and double KO (F) gonads. Only *Maf* KO gonads and mesonephroi contain bright GFP+ cells (C,D,F). (G-L) Immunofluorescent images of E13.5 XY CD-1 (G), E13.5 XX CD-1 (H), E12.5 XY C57BL/6 (I), E12.5 XY *Cx3cr1*-GFP (J), E12.5 XY CD-1 (K), and E13.5 XX CD-1 (L) gonads. G', H', and L' are higher-magnification images of the boxed regions in G, H, and L. CD11b-bright gonadal and mesonephric cells (arrowheads in G-K) do not express markers of gonadal and mesonephric macrophages (arrows in G-K) such as AIF1, NRP1, MRC1, *Cx3cr1*-GFP, and CSF1R. (L) CD11b-bright cells possess a U-shaped nucleus characteristic of monocytes. Thin scale bars, 50 μm. Thick scale bars, 12.5 μm.

**Supplemental Figure S6. MAFB and MAF are expressed in F4/80+ gonadal macrophages, but not in CD11b-bright monocytes.** Immunofluorescent images of E10.5 CD-1 urogenital ridge (A and B), E12.5 XY gonad/mesonephros (C-F), and E13.5 XY (G) gonads. A', B', and G' are higher-magnification images of the boxed regions in A, B, and G. Yellow dashed outlines in A and B mark gonad boundaries. (A and B) F4/80+ macrophages (arrowheads in A' and B') express MAFB and MAF, although there are occasional F4/80+ cells that do not express MAFB (arrow in A'). (C-G) F4/80+ macrophages (arrowheads in C-G) express both MAFB and MAF, while CD11b-bright monocytes (arrows in E and F) do not express MAFB or MAF. Note that MAFB is expressed in gonadal and mesonephric mesenchyme starting at E10.5 and MAF is expressed in gonadal mesenchyme starting at E12.0, consistent with our previous report [1]. Yellow dashed outline in G' indicates cord-interstitium boundary. (G) MAFB (as visualized by *Mafb*-GFP expression) and MAF expression are maintained in E13.5 gonadal macrophages (G', arrowheads). tc, testis cords; i, interstitium. Scale bars, 50 μm.

**Supplemental Figure S7. VEGFA treatment *ex vivo* disrupts vascular remodeling and hinders Leydig cell differentiation.** Immunofluorescent (A-D) and qRT-PCR (E) analyses of

48-hour ex vivo gonad culture of E12.5 CD-1 gonads (in 5% FBS media), showing that changes in vascular patterning induced by $VEGFA_{165}$ treatment (50 ng/mL) resulted in a decreased number of Leydig cells (*Cyp11a1*, *Hsd3b1*, and *Cyp17a1* mRNA and CYP11A1 immunostain). Note that $VEGFA_{165}$ administration also resulted in a reduction of germ cell numbers (*Ddx4*, *Kit*, and *Pou5f1* mRNA and DDX4 immunostain). Scale bars, 100 μm. All graph data are represented as mean +/- SD. *, *P*<0.05; **, *P*<0.01 (Student t-test).

**Supplemental Figure S8. Effects of vasculature and interstitial signaling pathways on Leydig cell differentiation.** (A-D) Immunofluorescence images of E12.5 CD-1 gonads cultured for 48 hours ex vivo with 10% FBS plus vehicle control (A), 50 ng/mL PDGF-BB (B), 1.8 μg/mL VEGFR-TKI II (C), or 50 ng/mL PDGF-BB plus 1.8 μg/mL VEGFR-TKI II (D), showing that PDGF-BB treatment has no impact on Leydig cell differentiation when vasculature is eliminated with VEGFR-TKI II. Scale bars, 100 μm. (E) qRT-PCR analyses for gonads cultured in the conditions shown in A-D. (F) qRT-PCR analyses of E13.5 XY *Mafb*-heterozygous;*Maf* KO gonads relative to controls, examining pathways required for Leydig cell differentiation, including Pdgf (*Pdgfa* and *Pdgfra*), desert hedgehog (*Dhh* and *Ptch1*), and Notch (*Hes1*, *Hey1*, and *Heyl*). All graph data are represented as mean +/- SD. ***, *P*<0.001; **, *P*<0.01 (Student t-test).

**Supplemental Table S1. Excel file of microarray gene expression data.** Excel file with worksheets containing data for all genes, upregulated genes, and downregulated genes in E12.5 XY *Mafb*-heterozyous;*Maf* KO ($Mafb^{GFP/+}$;$Maf^{-/-}$) GFP-expressing cells versus control

($Mafb^{GFP/+}$;$Maf^{+/-}$) GFP-expressing cells. Expression fold change data values have been transformed into log base 2.

**Supplemental Table S2. Primary antibodies used for immunofluorescence.**

| Primary Antibody | Dilution | Source/Reference | RRID |
|---|---|---|---|
| Rabbit anti-AIF1 (IBA1) | 1:1,000 | Wako #019-19741 | AB_839504 |
| Goat anti-AMH | 1:500 | Santa Cruz #sc-6886 | AB_649207 |
| Rat anti-CD11b | 1:250 | BD Biosciences #550282 | AB_393577 |
| Rat anti-CD45 | 1:300 | BioLegend #103101 | AB_312966 |
| Rat anti-CD45, Alexa488 | 1:300 | BioLegend #103121 | AB_493532 |
| Rat anti-CDH1 | 1:500 | Thermo Fisher #13-1900 | AB_2533005 |
| Rat anti-CDH5 | 1:500 | BD Biosciences #550548 | AB_2244723 |
| Rabbit anti-CSF1R | 1:1,000 | Santa Cruz #sc-692 | AB_631025 |
| Rabbit anti-CYP11A1 | 1:500 | D. Wilhelm [2] | N/A |
| Rabbit anti-CYP17A1 | 1:500 | Santa Cruz #sc-46081 | AB_2088659 |
| Rabbit anti-DDX4 (MVH) | 1:1,000 | Abcam #ab13840 | AB_443012 |
| Rat anti-F4/80 | 1:2,000 | Bio-Rad #MCA497RT | AB_1102558 |
| Goat anti-FOXL2 | 1:250 | Novus #NB100-1277 | AB_2106188 |
| Chicken anti-GFP | 1:1,000 | Aves #GFP-1020 | AB_10000240 |
| Rabbit anti-GFP | 1:1,000 | Thermo Fisher #A11122 | AB_221569 |
| Rabbit anti-HSD3B1 | 1:2,000 | K. Morohashi [3] | N/A |
| Rat anti-ICAM2 (CD102) | 1:1,000 | Bio-Rad #MCA2295 | AB_566421 |
| Rabbit anti-MAF | 1:2,000 | Bethyl #A300-613A | AB_2137521 |
| Rabbit anti-MAFB | 1:2,000 | Bethyl #A300-612A | AB_2265853 |
| Rabbit anti-MAFB | 1:500 | Millipore-Sigma #HPA005653 | AB_1079293 |
| Rat anti-MRC1 (CD206) | 1:1,000 | Bio-Rad #MCA2235T | AB_1101333 |
| Rabbit anti-NR5A1 | 1:2,000 | K. Morohashi [4] | N/A |
| Goat anti-NRP1 | 1:300 | R&D Systems #AF566 | AB_355445 |

| | | | |
|---|---|---|---|
| Goat anti-PECAM1 | 1:300 | R&D Systems #AF3628 | AB_2161028 |
| Rat anti-PECAM1 | 1:250 | BD Biosciences #553370 | AB_394816 |
| Rabbit anti-SOX9 | 1:4,000 | Millipore-Sigma #AB5535 | AB_2239761 |
| Rabbit anti-SYCP3 | 1:500 | Novus #NB300-231 | AB_10002746 |

**Supplemental Table S3. Sequences of primers used for qRT-PCR analyses.**

| **Gene name** | **Sequence (5' to 3')** |
|---|---|
| *Amh* forward | CCACACCTCTCTCCACTGGTA |
| *Amh* reverse | GGCACAAAGGTTCAGGGGG |
| *Arx* forward | CAAGGATGGTGAGGACAGC |
| *Arx* reverse | TCTGGAACCACACCTGGACT |
| *Cdh5* forward | TCCTCTGCATCCTCACTATCACA |
| *Cdh5* reverse | GTAAGTGACCAACTGCTCGTGAAT |
| *Cxcl12* forward | TGCATCAGTGACGGTAAACCA |
| *Cxcl12* reverse | TTCTTCAGCCGTGCAACAATC |
| *Cyp11a1* forward | TGGCCCCATTTACAGGGAGAA |
| *Cyp11a1* reverse | GGCATCTGAACTCTTAAACAGGA |
| *Cyp17a1* forward | CAGAGAAGTGCTCGTGAAGAAG |
| *Cyp17a1* reverse | AGGAGCTACTACTATCCGCAAA |
| *Ddx4* forward | TACTGTCAGACGCTCAACAGGA |
| *Ddx4* reverse | ATTCAACGTGTGCTTGCCCT |
| *Dhh* forward | CTTGGCACTCTTGGCACTATC |
| *Dhh* reverse | GACCCCCTTGTTACCCTCC |
| *Foxl2* forward | GCTACCCCGAGCCCGAAGAC |
| *Foxl2* reverse | GTGTTGTCCCGCCTCCCTTG |
| *Gapdh* forward | AGGTCGGTGTGAACGGATTTG |
| *Gapdh* reverse | TGTAGACCATGTAGTTGAGGTCA |
| *Hes1* forward | ATAGCTCCCGGCATTCCAAG |
| *Hes1* reverse | GCGCGGTATTTCCCCAACA |

| | |
|---|---|
| *Hey1* forward | GCGCGGACGAGAATGGAAA |
| *Hey1* reverse | TCAGGTGATCCACAGTCATCTG |
| *Heyl* forward | CAGCCCTTCGCAGATGCAA |
| *Heyl* reverse | CCAATCGTCGCAATTCAGAAAG |
| *Hsd3b1* forward | CAAGTGTGCCAGCCTTCATCT |
| *Hsd3b1* reverse | TTCATGATTCTGTTCCTCGTGG |
| *Jag1* forward | TGACATGGATAAACACCAGCA |
| *Jag1* reverse | GCAGCCCACTGTCTGCTATAC |
| *Kit* forward | CATGGCGTTCCTCGCCT |
| *Kit* reverse | GCCCGAAATCGCAAATCTTT |
| *Kitl* forward | TCTGCGGGAATCCTGTGACT |
| *Kitl* reverse | TGGAAGATTTGCCACCAGTTT |
| *Nestin* forward | GCTGGAACAGAGATTGGAAGG |
| *Nestin* reverse | CCAGGATCTGAGCGATCTGAC |
| *Nr2f2* forward | TCAACTGCCACTCGTACCTG |
| *Nr2f2* reverse | CCATGATGTTGTTAGGCTGCAT |
| *Pdgfa* forward | CAGTGTCAAGGTGGCCAAAGT |
| *Pdgfa* reverse | TGGTGTGGGTTCAGGTTGGA |
| *Pdgfra* forward | TCCATGCTAGACTCAGAAGTCA |
| *Pdgfra* reverse | TCCCGGTGGACACAATTTTTC |
| *Pou5f1* forward | GGAGGAAGCCGACAACAATGA |
| *Pou5f1* reverse | TCCACCTCACACGGTTCTCAA |
| *Ptch1* forward | AAAGAACTGCGGCAAGTTTTT |
| *Ptch1* reverse | CTTCTCCTATCTTCTGACGGGT |
| *Sox9* forward | GCGGAGCTCAGCAAGACTCTG |
| *Sox9* reverse | ATCGGGGTGGTCTTTCTTGTG |
| *Wnt4* forward | AGACGTGCGAGAAACTCAAAG |
| *Wnt4* reverse | GGAACTGGTATTGGCACTCCT |

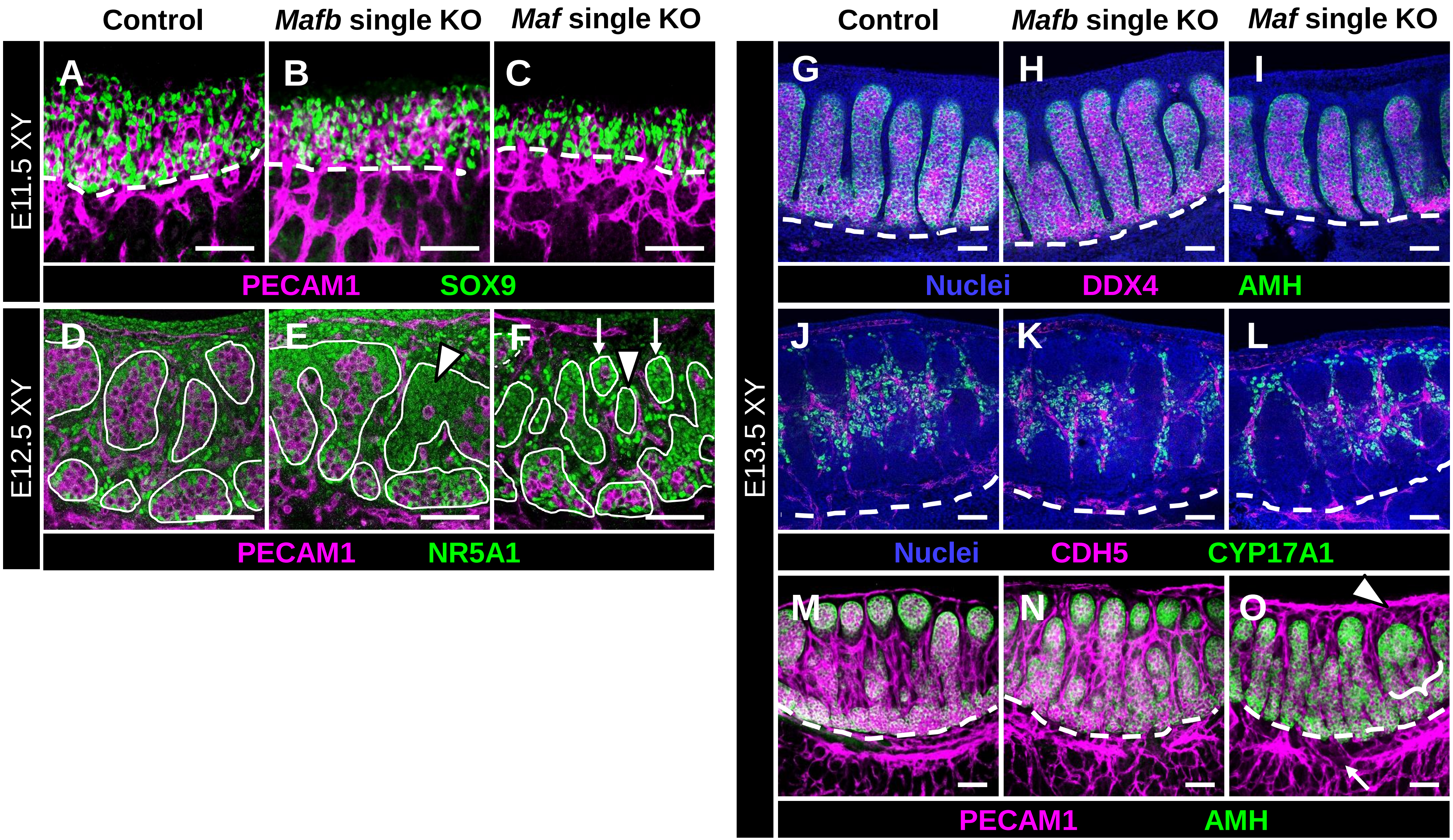


**Supplemental Figure S1**

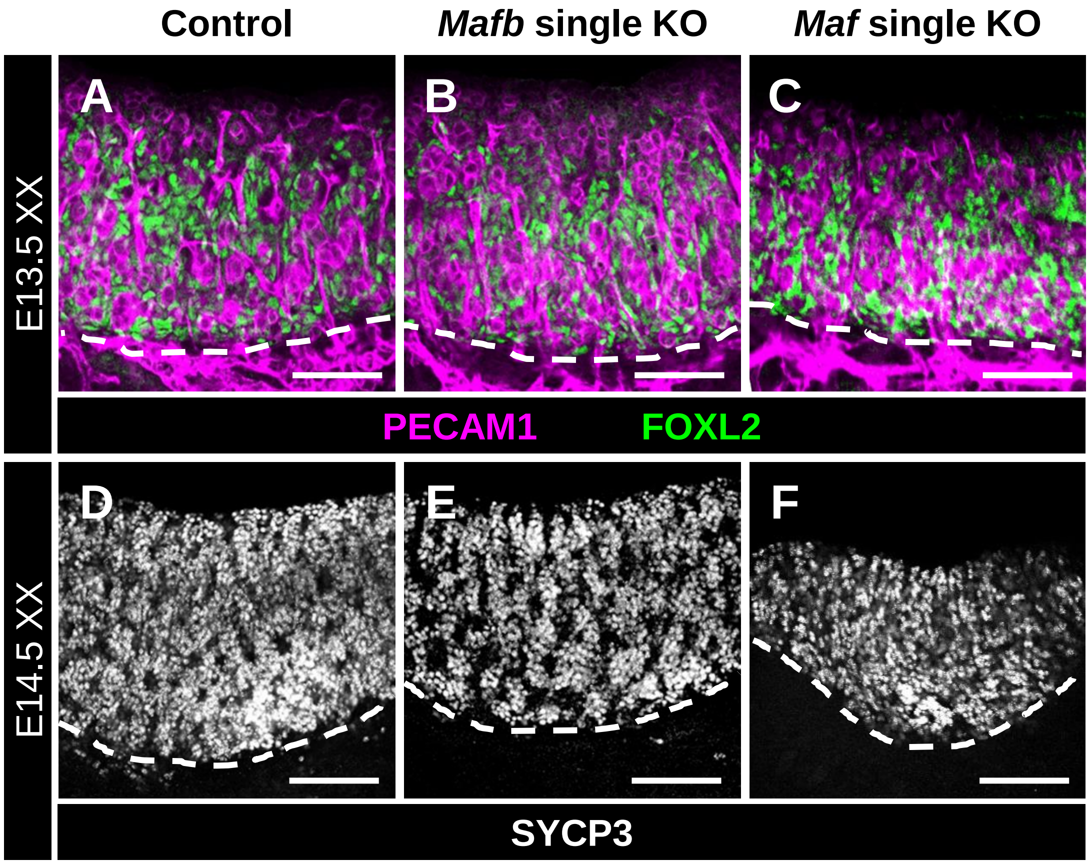


Supplemental Figure S2

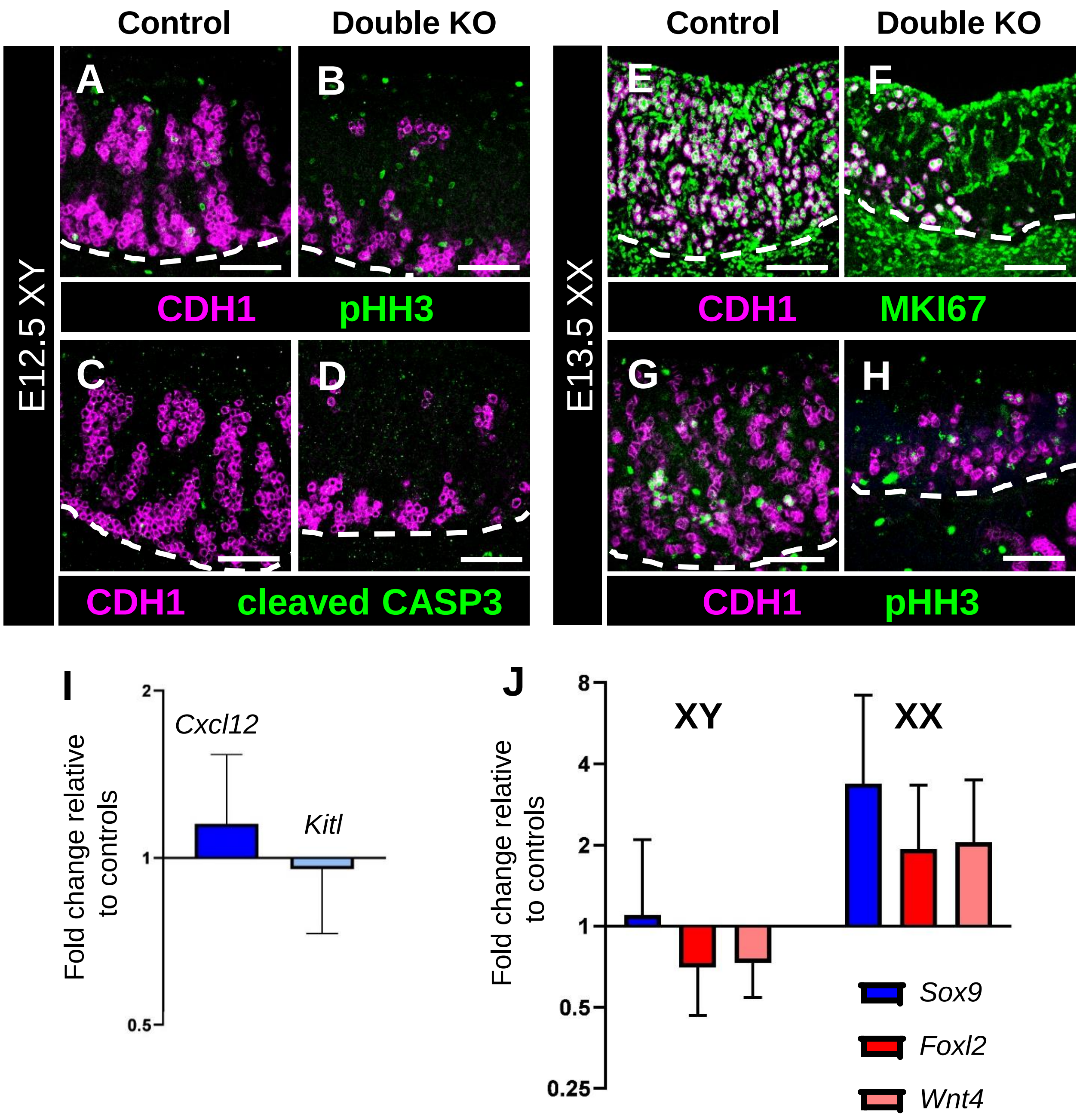


**Supplemental Figure S3**

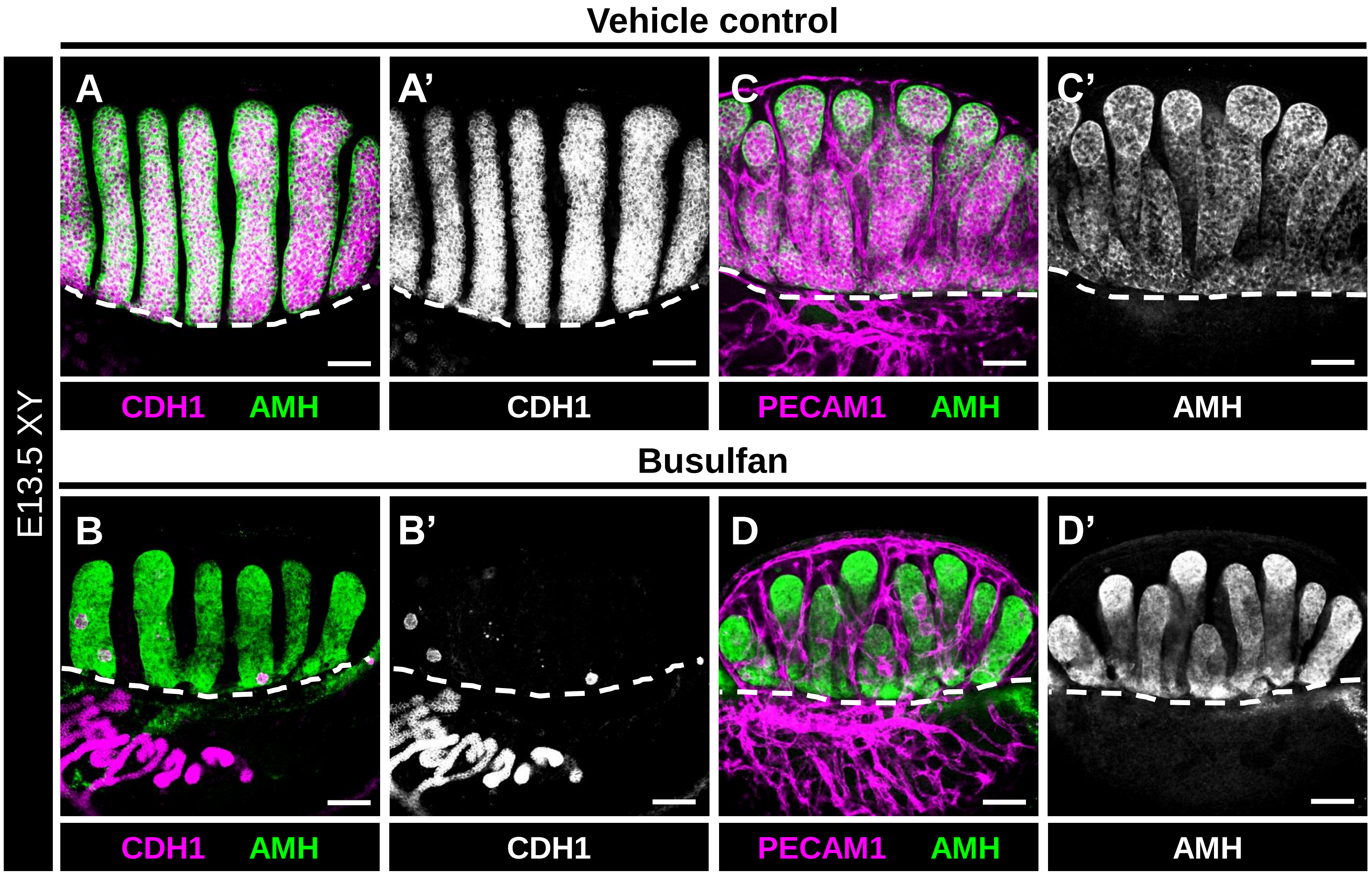


**Supplemental Figure S4**

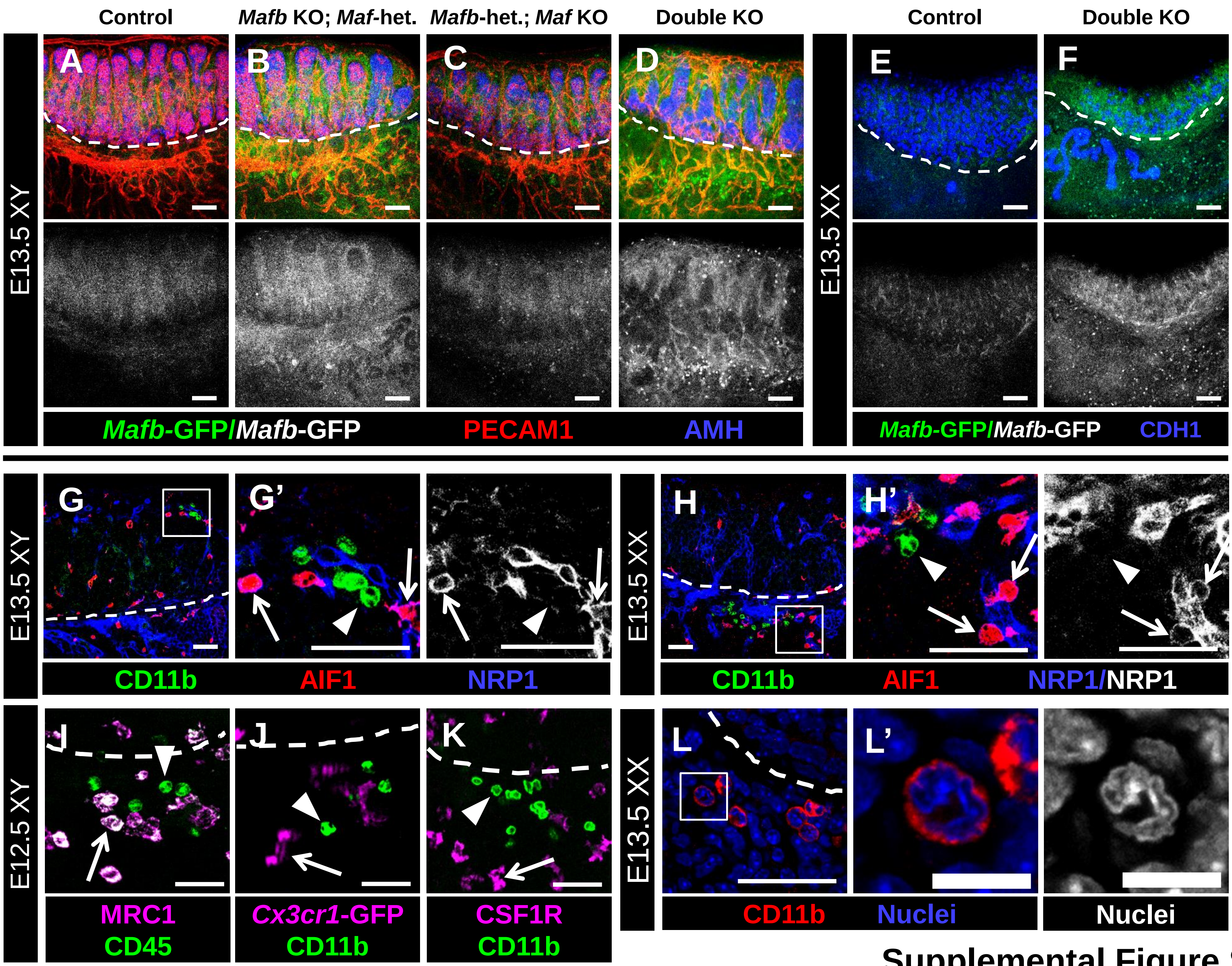


**Supplemental Figure S5**

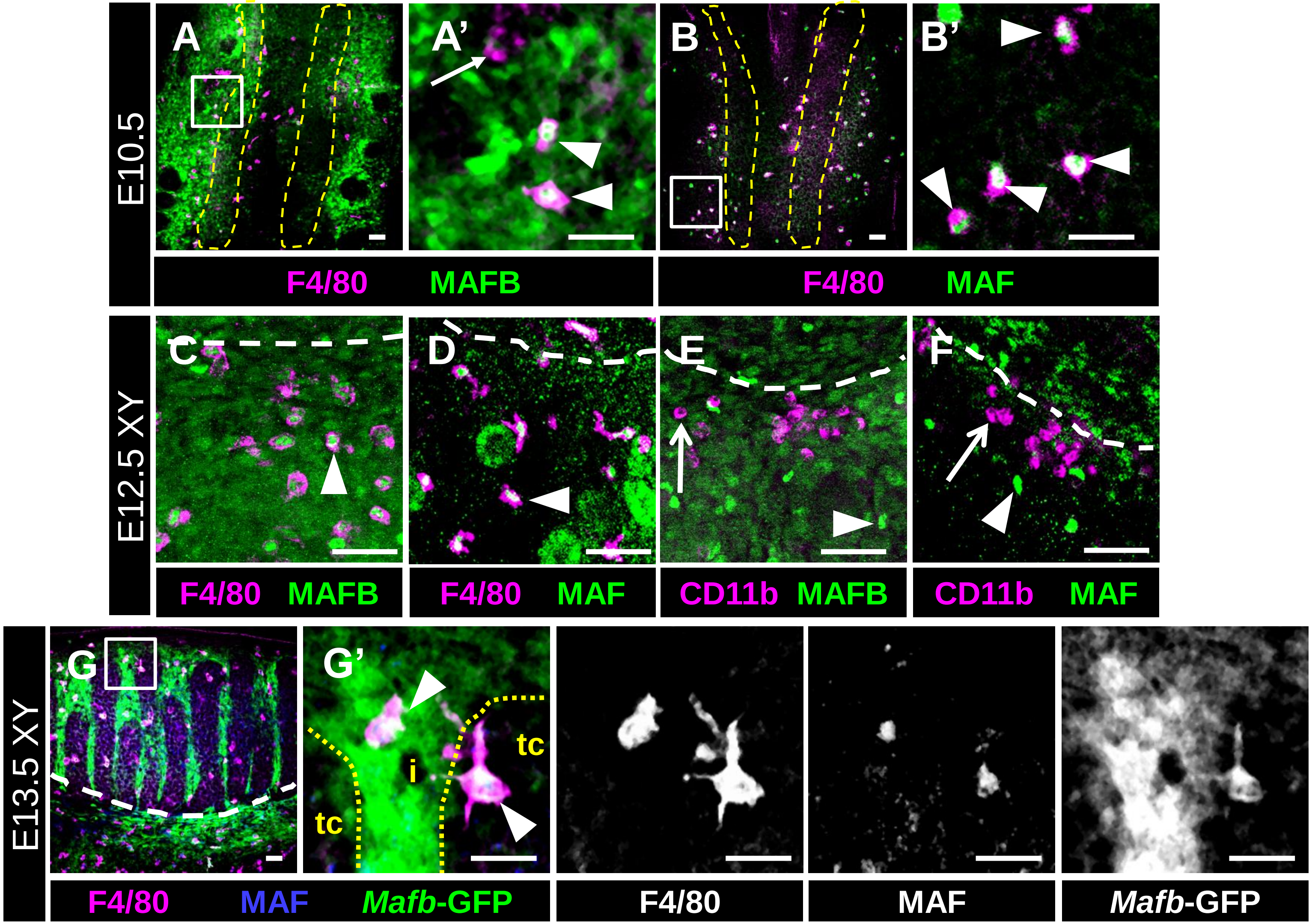


Supplemental Figure S6

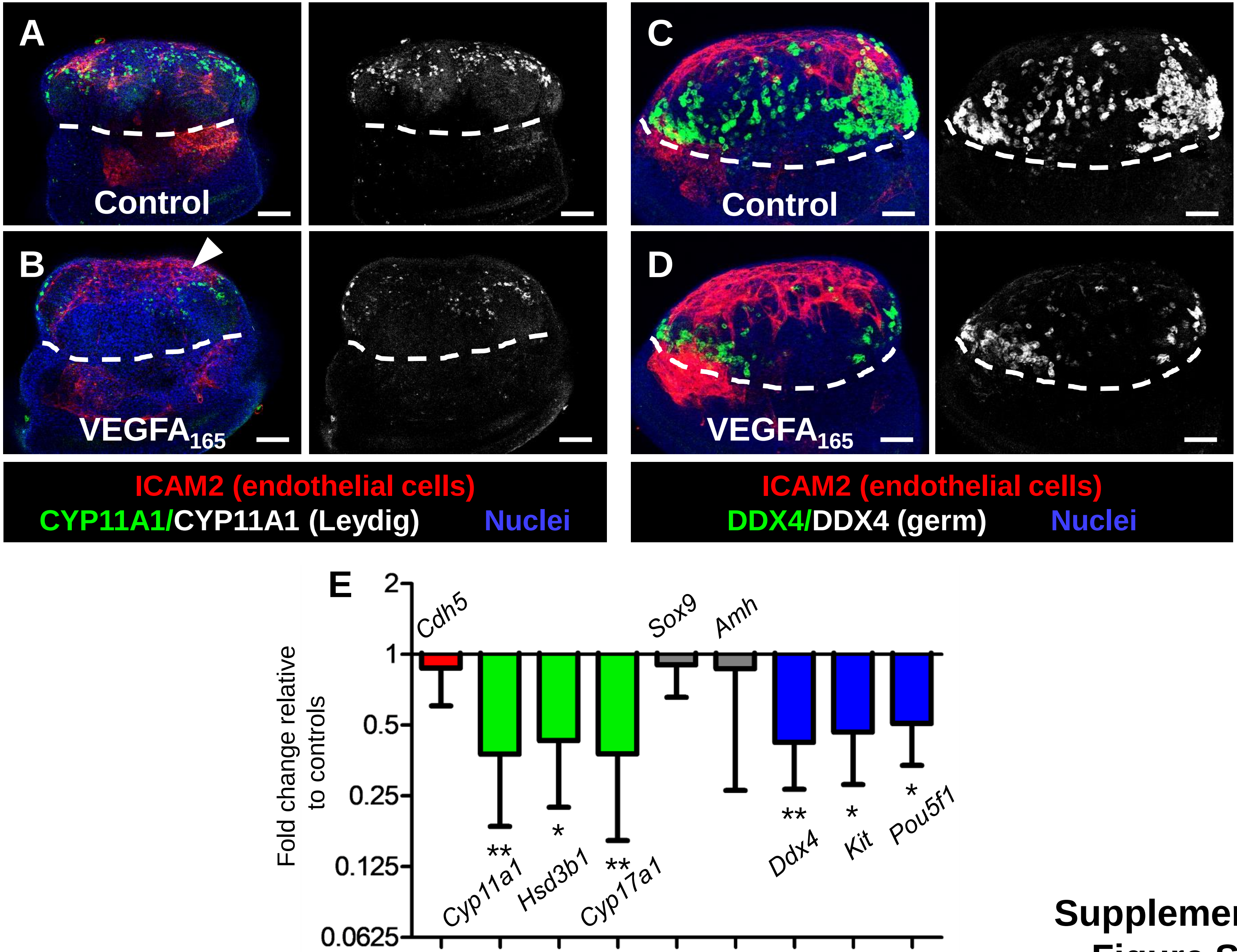


**Supplemental Figure S7**

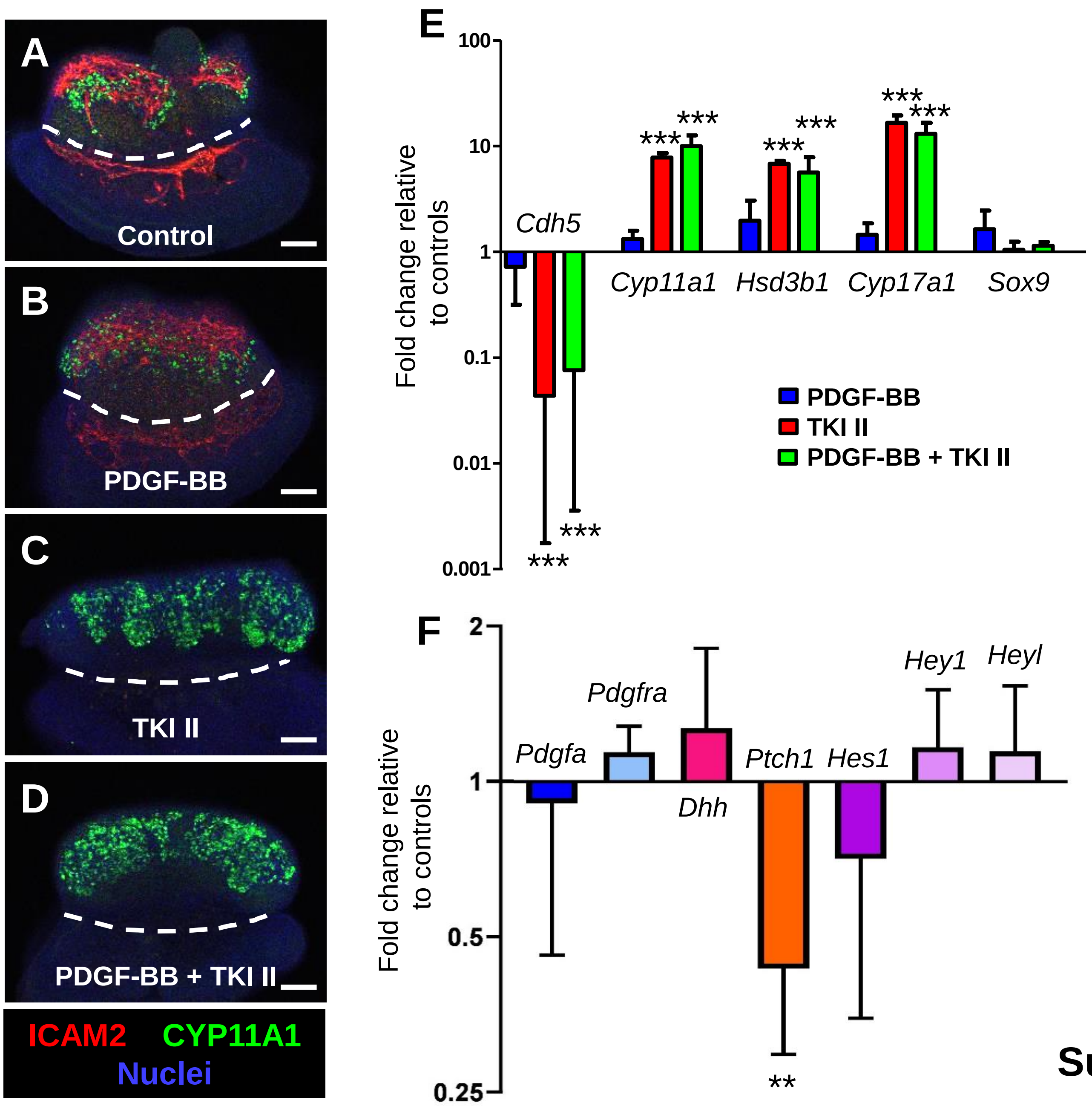


Supplemental Figure S8
